# TP53-mediated bidirectional lineage plasticity drives alveolar epithelial cell extrusion and tissue remodeling

**DOI:** 10.64898/2026.06.24.732965

**Authors:** Jeremy Morowitz, Thomas Whitlow, Naoya Miyashita, Khaliun Enkhbayar, Aditya Pratapa, Rohit Singh, Aleksandra Tata, Purushothama Rao Tata

## Abstract

Cell extrusion contributes to epithelial homeostasis, but its dysregulation can lead to tumorigenesis or degeneration. A fine balance in this process is therefore essential for tissue integrity. Yet the cell types and states vulnerable to extrusion, and the mechanisms that drive it, remain elusive. Here, using spatial maps of cell states in human idiopathic pulmonary fibrosis (IPF) we find that aberrant TP53 activation in alveolar epithelial cells drives cell extrusion. Genetic modulation of TP53 specifically in alveolar epithelial type 1 cells (AT1) was sufficient to induce plasticity and subsequent extrusion as demonstrated by lineage tracing and live imaging. Strikingly, single cell and bulk transcriptome profiling revealed aberrant TP53 drives AT1 cells to acquire a transitional state mirroring AT2-derived regeneration associated intermediate states. Critically, loss of AT1 derived transitional state triggers a compensatory AT2-derived regenerative response, establishing a bidirectional transitional state that activates myofibroblasts and remodels the alveolus. Together, our study implicates AT1 plasticity and their reversion as an unrecognized driver of epithelial cell loss and establishes bidirectional transitional state as a central mechanism underlying progression of fibrotic remodeling.

## Introduction

Cell extrusion is a conserved epithelial process that eliminates unfit, damaged, or supernumerary cells while preserving barrier integrity^1–3^. Dysregulation of this process can lead to opposing pathologies: insufficient extrusion permits the retention of aberrant cells and contributes to tumorigenesis, whereas excessive extrusion depletes essential cell populations and drives tissue degeneration. A fine balance in extrusion is therefore essential for tissue integrity. Yet the cell types and intermediate states that are vulnerable to extrusion, and the cues that trigger their loss, remain incompletely defined.

Cellular plasticity has emerged as a fundamental property of adult tissues, enabling differentiated cells to adapt to physiological stress, injury, and disease. Across mammalian organs, terminally differentiated cells can re-enter the cell cycle, dedifferentiate, or transit through intermediate cell states to drive tissue repair^4–6^. Beyond regeneration, such plasticity is increasingly recognized as a driver of pathological remodeling and tumorigenesis, in which cells of origin acquire altered identities that subvert normal tissue architecture^7,8^. Critically, the transient intermediate states adopted by plastic cells are often the very populations vulnerable to extrusion, coupling these two processes during epithelial remodeling and disease.

The lung exemplifies the need for tightly orchestrated cellular plasticity, balancing efficient gas exchange with maintenance of a durable epithelial barrier. The distal lung is organized into grape-like alveolar clusters lined by elegantly thin alveolar type 1 (AT1) cells, which mediate gas exchange, and cuboidal (alveolar type 2) AT2 cells, which secrete pulmonary surfactant and serve as facultative stem cells^9,10^. Upon injury and AT1 cell loss, AT2 cells differentiate to replace AT1 cells and self-renew to replenish the AT2 pool^9–14^. Recent work also established that this differentiation proceeds through a transient intermediate cell state, which we have termed the “Pre-AT1 Transitional State” (PATS)^15^, and which has been described in parallel as the *Krt8*+ Alveolar Differentiation Intermediate (ADI)^16^ and the Damage-Associated Transient Progenitor (DATP)^17^. The currently accepted paradigm therefore casts the alveolar epithelial continuum as unidirectional: AT2 stem cells transit through PATS into terminally differentiated AT1 cells^18–20^. In addition, recent studies also suggested that AT1 cells themselves exhibit plasticity, however, this is currently debate. Some studies proposed a bidirectional lineage relationship between AT1 and AT2 cells, in which AT1 cells could dedifferentiate into AT2 cells and self-renew^21^. Subsequent work demonstrated that modulation of YAP/TAZ signaling can drive reversion of AT1 cells toward an AT2-like state^22,23^, and that hyperoxia-induced stress similarly elicits AT1-to-AT2 plasticity^22^. In contrast, recently developed intersectional lineage-tracing strategies established that AT1 cells do not contribute to the AT2 pool under homeostatic conditions or after injury, but can acquire plasticity in the context of *Kras*-driven tumorigenesis^24^. AT1-specific *Egfr* mutation has further been shown to drive a histologically distinct subtype of lung adenocarcinoma^25,26^. In parallel, classical histopathological studies of pulmonary fibrosis described “desquamated epithelial cells with elongated morphology”^27^ adjacent to regions of interstitial thickening^28^, and modern single-cell analyses of human IPF^29–34^ and COVID-19-associated ARDS^35^ have further suggested an analogous population of detached, AT1-like cells of unresolved identity and origin. Additionally, studies have found that epithelial cell extrusion contributes to the denuding of airways in IPF^36^ as well as in the inflammatory-damage associated with bronchoconstriction in asthma^37^. Interestingly, extrusion has been demonstrated to preserve barrier function even in response to cellular overcrowding or mechanical constriction^1–3^, which is integrally related to disease of the lung. Despite these observations, we currently lack a clear mechanistic understanding of whether and how AT1 cells acquire plasticity in fibrotic lung diseases and contributes to extrusion and tissue pathology.

In this study, we set out to define the contribution of AT1 cell plasticity and extrusion to lung homeostasis and disease. Using spatial transcriptomic analysis of human IPF tissue, genetic modulation and lineage tracing specifically in AT1 cells, live imaging, and bulk and single-cell transcriptomics, we report that TP53-mediated AT1 plasticity drives their transition into a PATS-like cell state, their extrusion from the alveolar epithelium. This that collectively leads to remodeling of the alveolar compartment with implications to distal lung diseases including fibrosis.

## Results

### Epithelial detachment in human idiopathic pulmonary fibrosis is enriched for AT1 cells with elevated TP53 signaling

A hallmark of human fibrotic lungs is thickening of the alveolar septal walls, loss of alveolar epithelial cells, and the emergence of aberrant alveolar epithelial cell states (Figure 1a, Supplementary Figure 1a). To characterize the cellular features of this remodeling, we performed immunofluorescent analysis of human fibrotic lung tissue, which revealed profound disruption of the alveolar architecture that progressively propagates into neighboring regions. Notably, the epithelium adjacent to thickened septal walls was frequently observed to detach from the underlying basement membrane (Figure 1b). These detached cells expressed SFN, a canonical marker of PATS, while additional populations of detached cells expressed AGER and SFTPC, markers of AT1 and AT2 cells, respectively (Figure 1b). Such epithelial detachment was apparent both as multicellular sheets and as individual AT1 cells (Supplementary Figure 1b).

**Figure 1:**
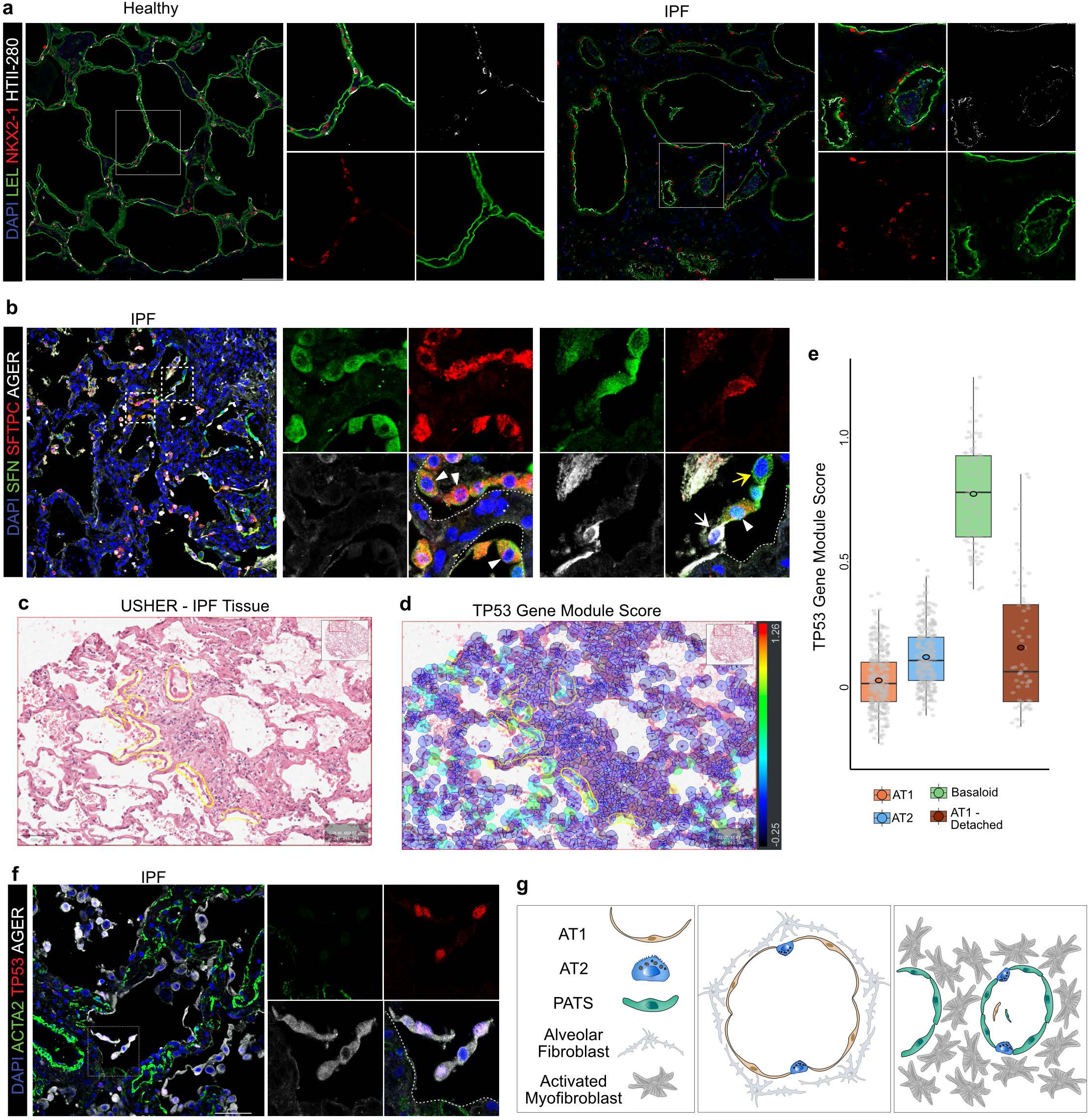
Alveolar epithelial dysregulation in human fibrotic lung tissue is characterized by elevated TP53 signaling. a. Immunostaining for LEL, NKX2-1, and HTII-280 in human lung tissue, representative images healthy and IPF lung. DAPI stains nuclei. White box indicates region of single-channel images. Scale bar: 100 µm. b. Immunostaining for SFN, SFTPC, and AGER in human IPF lung tissue. The white dotted line represents the edge of the basement membrane, with white triangles indicate regions of epithelium that have detached. White arrow indicates AGER+ cell detaching, while the yellow arrow indicates an SFN cell detaching. DAPI stains nuclei. White box indicates region of single-channel images. Scale bar: 100 µm. c. USHER analysis of regions of human IPF tissue that exhibit epithelial detachment. d. TP53 signaling signatures, comparing regions of detached cells with AT1 morphology to other alveolar epithelium classified as “AT1”, “AT2”, and cells classified as “Basaloid”, based on gene module scores. e) USHER analysis of regions of human tissue that exhibit epithelial detachment, showing visual representation of TP53 gene module score. f) Immunostaining for ACTA2, TP53, and AGER in IPF lung The white dotted line represents the edge of the basement membrane. DAPI stains nuclei. White box indicates region of single-channel images. Scale bar: 100 µm. g) Schematic illustrating the epithelial (AT1, AT2, and PATS), and mesenchymal (alveolar fibroblasts and activated myofibroblasts) cells organization in healthy and IPF lung.

To dissect the molecular mechanisms underlying this epithelial detachment and to define the contribution of distinct alveolar cell types, we employed USHER, a newly developed spatial transcriptomics analysis tool^38^. Briefly, USHER integrates rich single-cell RNA-sequencing (scRNA-seq) data with sparse spatial transcriptomic measurements (Xenium) and uses the scRNA-seq reference to impute gene expression onto the spatial coordinates. This enrichment of the Xenium readout enables robust cell type classification and high-resolution transcriptomic analysis directly on the underlying H&E image. We applied this method to a human IPF biopsy that contained clear regions of epithelial detachment. Although the detached cells displayed heterogeneous morphologies, distinct regions of long, thin, and flat AT1-like cells detaching from the basement membrane were readily apparent (Figure 1c). Given the prior implication of TP53 in alveolar epithelial cell-state regulation and its observed expression in IPF tissue, we used USHER to selectively interrogate these AT1-detached regions for TP53 signaling, scoring cells with a TP53 gene module comprising *SFN*, *GDF15*, *BAX*, *BBC3*, and *RRM2B*. Strikingly, regions of detached AT1 epithelial cells were strongly enriched for the TP53 module score (Figure 1d). Furthermore, direct comparison across cell types within the same biopsy revealed that the TP53 module score in detached epithelial regions was higher than that of resident AT1 and AT2 cells but lower than that of the aberrant basaloid population (Figure 1e). Together, these data suggest that TP53 signaling is mechanistically associated with AT1 cell detachment in human IPF tissue.

To further validate this association, we co-stained human IPF tissue for TP53 and the canonical AT1 marker AGER. Notably, in fibrotic regions marked by ACTA2 expression, AT1 cells that had detached from the basement membrane were positive for TP53 (Figure 1f, Supplementary Figure 1b). Taken together, our USHER-based spatial transcriptomic analysis and orthogonal immunofluorescent validation reveal that elevated TP53 signaling is a defining feature of epithelial cell detachment in human IPF and specifically implicate AT1 cells as active contributors to the epithelial cell loss and alveolar remodeling that are central to IPF pathology (Figure 1g).

### Stabilization of TP53 in AT1 cells leads to morphological changes and their lineage reversion *in vivo*

To test whether stabilized TP53 in AT1 cells recapitulates the phenotypes we observed in diseased human lungs, we established an AT1-specific mouse model of TP53 stabilization. For this, we crossed *Ager-CreERT2;R26R-tdTomato* (hereafter referred to as *Ager-tdT*) mice to *Mdm2* floxed animals to generate *Ager-CreERT2;R26R-tdTomato;Mdm2*^fl/fl^ mice (hereafter referred to as *Ager-Mdm2-KO*). In this model, we intraperitoneally administered tamoxifen on three consecutive days to label AT1 cells with tdTomato and concomitantly delete *Mdm2*, leading to stabilized TP53 expression specifically in AT1 cells (Figure 2a). To assess how stabilized TP53 affects AT1 cell identity and morphology *in vivo*, we collected lungs at days 7, 14, and 21 post-tamoxifen. First, we co-stained lung sections for canonical epithelial identity markers: HOPX for AT1 cells and DCLAMP for AT2 cells. Notably, tdTomato^+^ lineage-labeled AT1 cells in *Ager-Mdm2-KO* lungs displayed an aberrant, and rounded morphology, in stark contrast to the characteristic thin and squamous shape of homeostatic AT1 cells. Strikingly, a substantial fraction of tdTomato^+^ lineage-labeled cells expressed low levels of the canonical AT2 markers DCLAMP and SFTPC at all time points examined (Figure 2b, Supplementary Figure 2a-c).

**Figure 2:**
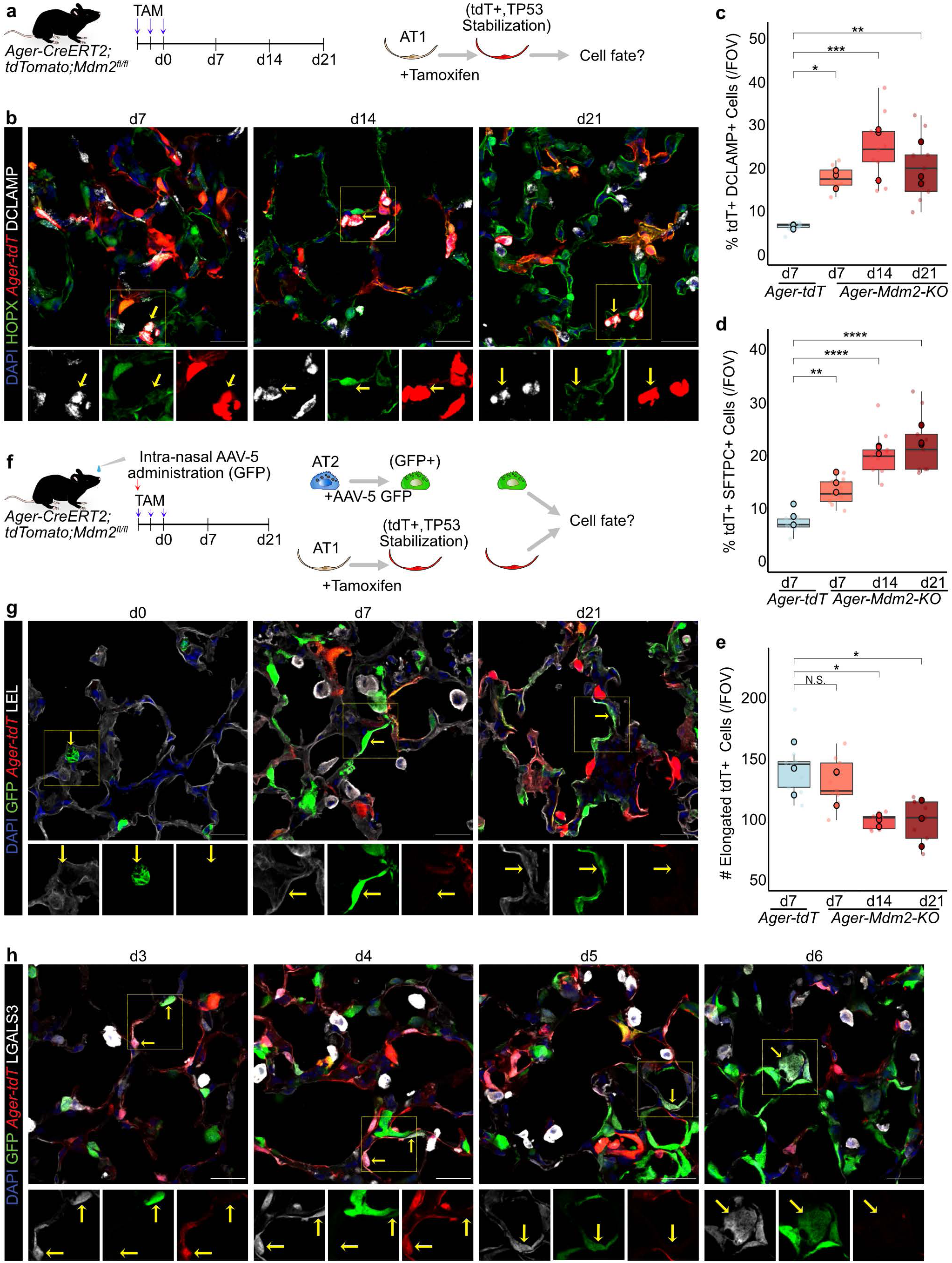
TP53 stabilization in AT1 cells drives cell identity dysregulation, AT1 cell loss, and compensatory AT2-driven regeneration. a. Experimental workflow for tamoxifen administration to lineage label and delete *Mdm2* in AT1 cells follow by tissue collection at days 7, 14, and 21. b. Immunostaining for HOPX, Ager-tdT, and DCLAMP in *Ager-Mdm2-KO* at days 7, 14, and 21. Scale bar: 25 µm. c. Quantification of the percentage of tdT^+^ lineage-labeled cells per field of view that also express DCLAMP. Larger opaque points represent biological replicates; smaller transparent points represent technical replicates. n=3. Data are presented as mean ± SEM. One-way ANOVA. *p = 0.0336, **p = 0.00207, *p = 0.0122. d. Quantification of the percentage of tdT^+^ lineage-labeled cells per field of view that express SFTPC. Larger opaque points represent biological replicates, smaller transparent points represent technical replicates. n=3. Data are presented as mean ± SEM. One-way ANOVA. **p = 0.00636, ***p = 2.66e-05, ***p = 5.67e-06. e. Quantification of number of elongated tdT^+^ lineage-labeled cells per field of view. Larger opaque points represent biological replicates, smaller transparent points represent technical replicates. n=3. Data are presented as mean ± SEM. One-way ANOVA. N.S. = not significant (p = 0.709), *p = 0.0339, *p = 0.0305. f. Schematic illustrating bidirectional lineage label model, with lineage labeling and *Mdm2* deletion in AT1 cells with three doses of tamoxifen, and lineage labeling of AT2 cells through intranasal administration of AAV-5 GFP. Tissue collection at days 0, 7, and 21. g. Immunostaining for GFP, Ager-tdT, and LEL in bi-directional lineage label model at days 0, 7, and 21. Scale bar = 25 µm. Images presented as z-projection of 2 stacks over 2 µm. h. Immunostaining for GFP, Ager-tdT, and LGALS3 in bi-directional lineage label model at days 3, 4, 5, and 6. Arrows indicate LGALS3+ PATS cells, seen as both GFP+ and tdT^+^, representing AT2- and AT1-derived PATS, respectively. Scale bar: 25 µm.

Quantification of these observations confirmed a significant number of cells expressing AT2 markers in AT1-lineage-labeled cells upon TP53 stabilization. In control *Ager-tdT* lungs, only about 6.38% of tdTomato^+^ lineage-labeled cells expressed DCLAMP, whereas in *Ager-Mdm2-KO* lungs this fraction rose to about 17.59%, 24.68%, and 20.08% at days 7, 14, and 21, respectively (Figure 2c). Similarly, SFTPC was detected in about 6.85% of tdTomato^+^ cells in controls compared to about 13.01%, 20.28%, and 22.05% at days 7, 14, and 21 in *Ager-Mdm2-KO* lungs (Figure 2d). Together, these data indicate that stabilization of TP53 in AT1 cells drives both morphological alterations and loss of AT1 cell identity *in vivo*. As expected, TP53 stabilized cells did not show markers of replication, as revealed by immunostaining for Ki67, suggesting that the gradual increase in tdTomato+ cells is due to AT1 lineage reversion without their amplification (Supplementary Figure 2b, 2c). Of note, while quantifying AT2 marker expression in lineage-labeled cells, we noticed that *Ager-Mdm2-KO* lungs showed a substantial decrease in single-positive tdTomato^+^ cells with characteristic elongated AT1 cell morphology compared to controls (Figure 2e).

To rule out the possibility that the emergence of AT2 markers within the tdTomato^+^ population reflected leaky activity of the *Ager-CreERT2* driver in AT2 cells, we performed a complementary bidirectional lineage-tracing experiment. For this, we infected *Ager-Mdm2-KO* mice with AAV5-GFP, which selectively labels AT2 cells (Figure 2f). This dual-labeling strategy allowed us to unambiguously distinguish AT1-derived (tdTomato^+^) cells from AT2-derived (GFP^+^) cells. Lungs collected at day 0, 7, and 21 revealed that GFP^+^ AT2 cells comprised a population distinct from tdTomato^+^ lineage-labeled AT1 cells (Figure 2g). Interestingly, as early as day 7 we observed GFP^+^ cells with elongated and stretched morphology characteristic of AT2 to AT1 differentiation, and by day 7 we identified thin LEL^+^ AT1 cells that co-expressed GFP, indicating that AT2 cells had differentiated into AT1 cells in *Ager-Mdm2-KO* lungs. Consistent with this, lungs at earlier time points showed that GFP^+^ cells expressed markers of alveolar epithelial transitional state (here after referred to as PATS) as early as day 3 and persisting through day 6 (Figure 2h) Taken together, these data demonstrate that stabilization of TP53 in AT1 cells leads to AT1 cell loss and elicits a regenerative response in which AT2 cells differentiate via the PATS transitional state to replenish the lost AT1 cell pool.

### Stabilized TP53 in AT1 cells triggers their eventual loss through extrusion

Our observation that tdTomato^+^ lineage-labeled AT1 cells diminish while AAV5-GFP-labeled AT2 cells differentiate into AT1 cells in *Ager-Mdm2-KO* lungs prompted us to investigate the fate of TP53-stabilized AT1 cells. To this end, we assessed lung sections from days 7, 14, and 21 for evidence of AT1 cell loss (Figure 3a). We identified tdTomato^+^ lineage-labeled cells with a rounded morphology, detached from the alveolar epithelium and retaining strong TP53 expression (Figure 3b). To assess the frequency of this phenomenon in an unbiased manner, we collected bronchoalveolar lavage (BAL) fluid from control and *Ager-Mdm2-KO* mice at day 7 (n = 3 per group). The percentage of tdTomato^+^ lineage-labeled cells recovered in BAL fluid was significantly higher in *Ager-Mdm2-KO* mice compared to controls (Figure 3c, 3d). Together, these data corroborate that stabilization of TP53 in AT1 cells leads to their detachment and eventual loss from the alveolar epithelium.

**Figure 3:**
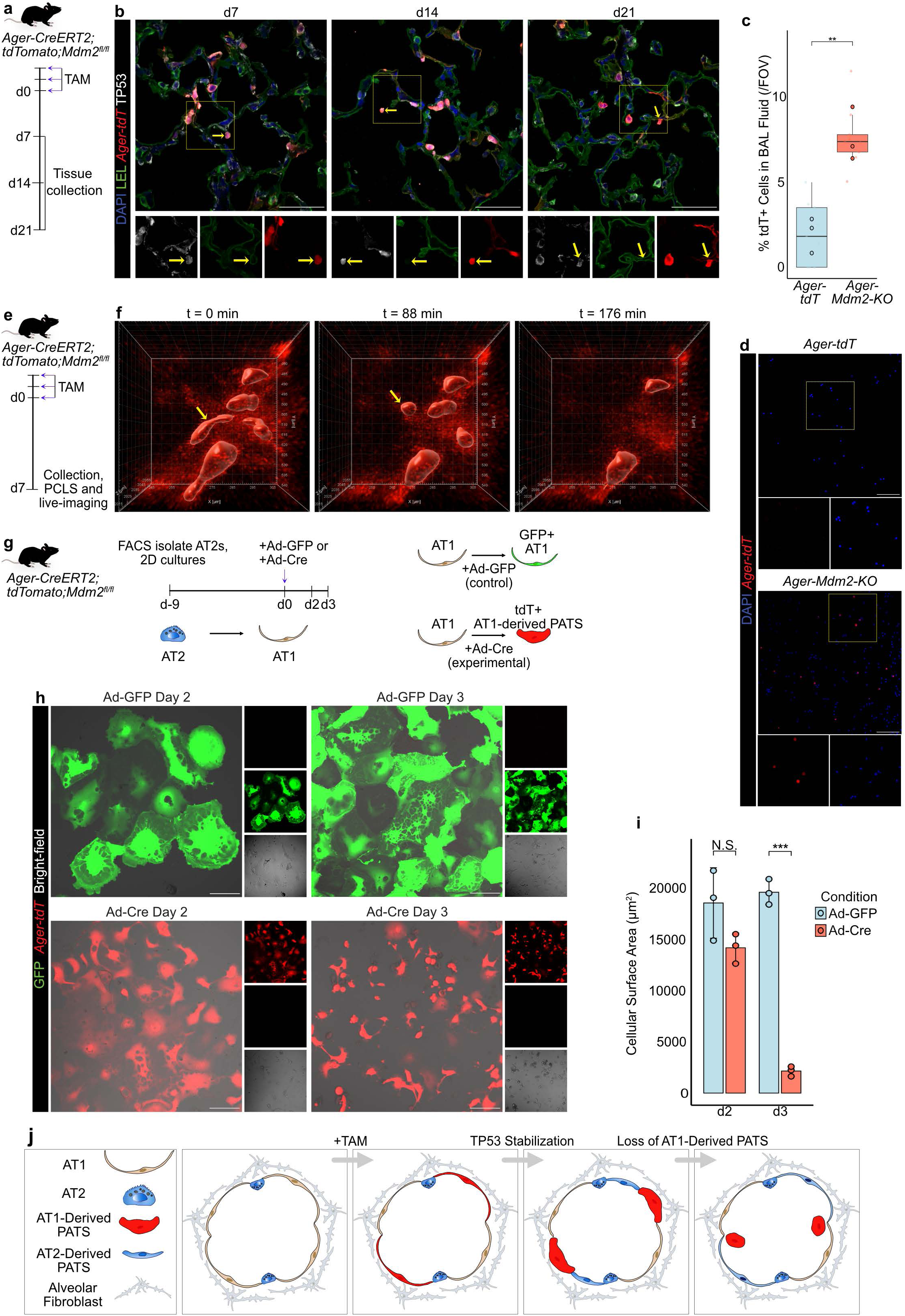
TP53 stabilization in AT1 cells induces a bidirectional PATS originating from both AT1 and AT2 cells. a. Schematic illustrating lineage labeling and *Mdm2* deletion in AT1 cells with three doses of tamoxifen, with tissue collection at days 7, 14, and 21. b. Immunostaining for LEL, Ager-tdT, and TP53 at days 7, 14, and 21. DAPI stains nuclei. Yellow arrow depicts tdT+TP53+ excluded cell. Yellow box indicates region of single-channel images. Scale bar: 50 µm. c. Quantification of the percentage of tdT^+^ lineage-labeled cells per field of view in BAL fluid. Larger opaque points represent biological replicates, smaller transparent points represent technical replicates. n=3. Unpaired two-tailed Welch’s t-test. **p = 0.0098. d. Images of tdT^+^ cells from *Ager-tdT* and *Ager-Mdm2-KO* BAL fluid collected at day 7. DAPI stains nuclei. Yellow box indicates region of single-channel images. Scale bar: 100 µm. e. Schematic illustrating lineage labeling and *Mdm2* deletion in AT1 cells with three doses of tamoxifen, with tissue collection at day f. Time frames from *Ager-Mdm2-KO* PCLS ex vivo cultures. Imaris software was used to apply cell mask outlines. Still frames represent lineage-labeled AT1 cells at t=0 min, t=88 min, and t=176 min, demonstrating loss of lineage-labeled AT1 cells. Yellow arrow indicates tdT^+^ cell undergoing extrusion. g. Schematic illustrating 2D culture experiments. AT2 cells were purified from *Ager- Mdm2-KO* mice and differentiated into AT1 cells on fibronectin for 9 days follow by administration of Ad-GFP (control) or Ad-Cre (experimental) virus. Imaged at days 2 and 3 of culture. h. Images showing GFP, tdT and bright field in 2D cultures of AT1 cells infected with indicated viruses. Scale bars: 100 µm. i. Quantification of cellular surface area from AT1 2D cultures infected with Ad-GFP (control) and Ad-Cre (experimental). n=3. Paired two-sample t-test. N.S. = not significant (p = 0.0626), **p = 0.00282. j. Schematic illustrating the alveolar cell types, highlighting that PATS can be derived from AT1 as well as AT2 cellular origins, and AT1-derived PATS induced by TP53 stabilization are lost, and further induce AT2-derived PATS.

To capture this loss in real time, we performed live imaging of precision-cut lung slices (PCLS) prepared from *Ager-Mdm2-KO* mice at day 7. For this, lungs were inflated with 1.5% low-melting agarose and sectioned at 750 µm on a compresstome. Lung slices were stained with LEL to mark the AT1 cell surface and imaged for 12 hours at 4-min intervals (Figure 3e and Supplementary Video 1 and 2). Lineage-labeled AT1 cells were initially observed as thin, squamous cells forming a continuous epithelial layer that lined the alveolar cup. However, over the course of hours, this morphology was progressively disrupted: tdTomato^+^ cells rounded up and were extruded from the alveolar cup into the luminal space (Figure 3f). Together, these live-imaging data reveal that stabilization of TP53 in AT1 cells drives morphological dysregulation followed by extrusion-mediated cell loss (Supplementary Figure 3c).

To dissect this process at higher resolution, we recapitulated TP53 stabilization in AT1 cells *in vitro*. For this, we isolated AT2 cells from *Ager-Mdm2-KO* mice by FACS and differentiated them into AT1 cells in monolayer cultures as described previously^39^. Upon spreading and acquisition of AT1 cell morphology, cultures were transduced with Adeno-Cre to simultaneously delete *Mdm2* (to stabilize TP53) and induce tdTomato lineage labeling. Parallel wells were transduced with Adeno-GFP as a control (Figure 3g). Consistent with our *in vivo* findings, Adeno-Cre-transduced cultures exhibited tdTomato expression accompanied by overt morphological changes in lineage-labeled AT1 cells (Figure 3h). Specifically, tdTomato^+^ AT1 cells rounded up and lost the characteristic thick microtubule bundles that define well-differentiated AT1 cells in a 2D monolayer culture. Quantification of these cultures revealed that AT1 cells with stabilized TP53 covered significantly less surface area than GFP-transduced controls (Figure 3i). Therefore, our *in vivo*, *in vitro*, and *ex vivo* data converge on the conclusion that stabilization of TP53 in AT1 cells reverts them towards PATS with accompanying morphological changes and their delamination from the alveolar epithelium. This in turn induces PATS from AT2 cells to replenish lost AT1 cells. Taken together, these data suggest that PATS emerge in two waves: one from reversion of AT1 cells (induced by TP53 stabilization), and another from AT2 cells (to replenish lost AT1 cells) (Figure 3j).

### Emergence of bidirectional PATS transiently induces myofibroblasts in alveolar niches

Previous studies revealed that emergence of PATS induces fibroblast-to-myofibroblast conversion during injury repair and in disease ^40^. Therefore, we sought to test whether a similar mesenchymal response accompanies this model. To this end, we co-stained lung sections from *Ager-Mdm2-KO* mice at days 7, 14, and 21 for galectin 3 (LGALS3), a canonical PATS marker, and alpha smooth muscle actin (ACTA2, also known as aSMA), a marker of myofibroblasts (Figure 4a). Notably, as early as day 7 we detected strong ACTA2 expression in the interstitial space underlying the epithelium of the alveolar cups, alongside tdTomato^+^ lineage-labeled cells that co-expressed LGALS3 (Figure 4b). This concurrent emergence of AT1-derived PATS cells and activated myofibroblasts as early as day 4 and persisted at day 14 and 21 (Figure 4b and Supplementary Figure 4a). Quantification of tissue coverage as a percentage of field of view further confirmed fibroblast to myofibroblast transition on day 7 in *Ager-Mdm2-KO* mice compared to controls (Figure 4c). Taken together, these data demonstrate that stabilization of TP53 in AT1 cells drives a coordinated sequence of epithelial cell loss, emergence of bidirectional PATS, myofibroblast activation, and progressive remodeling of the alveolar architecture (Figure 4d).

**Figure 4:**
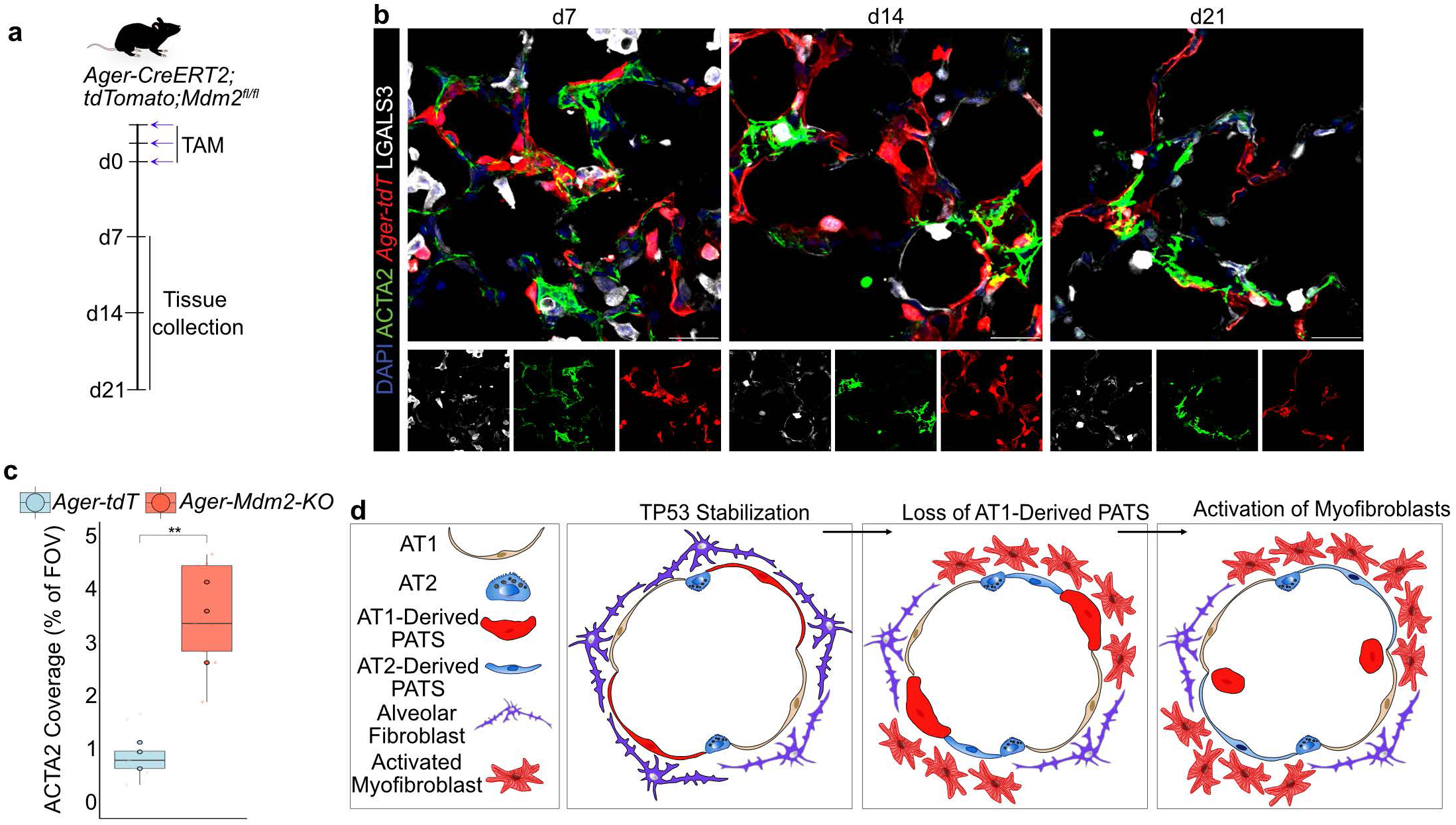
”Bi-directional” PATS induces conversion of fibroblasts to myofibroblast and tissue remodeling. a. Schematic illustrating lineage labeling and *Mdm2* deletion in AT1 cells with three doses of tamoxifen, with tissue collection at days 7, 14, and 21. b. Immunostaining for ACTA2, Ager-tdT, and LGALS3 at days 7, 14, and 21. DAPI stains nuclei. Scale bar: 25 µm. c. Quantification of ACTA2 coverage in *Ager-tdT* and *Ager-Mdm2-KO* lung sections collected at day 7, shown as a percentage of total field of view. Large opaque points represent biological replicates, n=3. Welch’s two-sample t-test. *p = 0.0205. d. Schematic illustrating the epithelial (AT1, AT2, and PATS) and mesenchymal (alveolar fibroblasts and activated myofibroblasts) cell organization in healthy lung tissue and fibrotic disease. This also shows TP53 stabilization, which leads to AT1-derived PATS, AT2-derived PATS, and subsequent fibroblast to myofibroblast activation, and alveolar remodeling.

### Bulk and single-cell transcriptomics reveal that TP53 stabilization drives AT1 and AT2 cells toward a convergent PATS state

Our observations in human tissue, mouse models, and *in vitro* cultures collectively indicate that stabilization of TP53 in AT1 cells reprograms alveolar epithelial cell identity and directs AT1 cells into a PATS-like state. To define this AT1-derived PATS state at the transcriptional level and compare it to previously characterized AT2-derived PATS, we performed bulk RNA sequencing on FACS-isolated tdTomato^+^ cells from four mouse models: (1) control AT1 cells from *Ager-CreERT2;R26R-tdTomato* (*Ager-tdT*); (2) AT1 with stabilized TP53 from *Ager-CreERT2;R26R-tdTomato;Mdm2*^fl/fl^ (*Ager-Mdm2-KO*); (3) control AT2 cells from *Sftpc-CreERT2;R26R-tdTomato* (*Sftpc-tdT*); and (4) AT2 with stabilized TP53 from *Sftpc-CreERT2;R26R-tdTomato;Mdm2*^fl/fl^ (*Sftpc-Mdm2-KO*) (Figure 5a). Differential expression analysis (as revealed by both heatmap and principal component analysis (PCA)) revealed distinct transcriptomic signatures across the four groups: *Ager-tdT* and *Sftpc-tdT* controls were the most divergent, whereas *Ager-Mdm2-KO* and *Sftpc-Mdm2-KO* samples converged toward each other and clustered between the two control groups (Figure 5b, 5c). Comparisons of *Sftpc-Mdm2-KO* versus *Sftpc-tdT* and of *Ager-Mdm2-KO* versus *Ager-tdT* revealed that *Mdm2* deletion and the subsequent TP53 stabilization induced upregulation of canonical PATS genes including *Sfn*, *Cldn4*, *Krt8*, and *Lgals3* in both cell-of-origin contexts (Supplementary Figure 5a, 5b). Volcano plots further highlighted additional differentially expressed genes between *Mdm2*^fl/fl^ and control samples in each comparison (Figure 5d, 5e).

**Figure 5:**
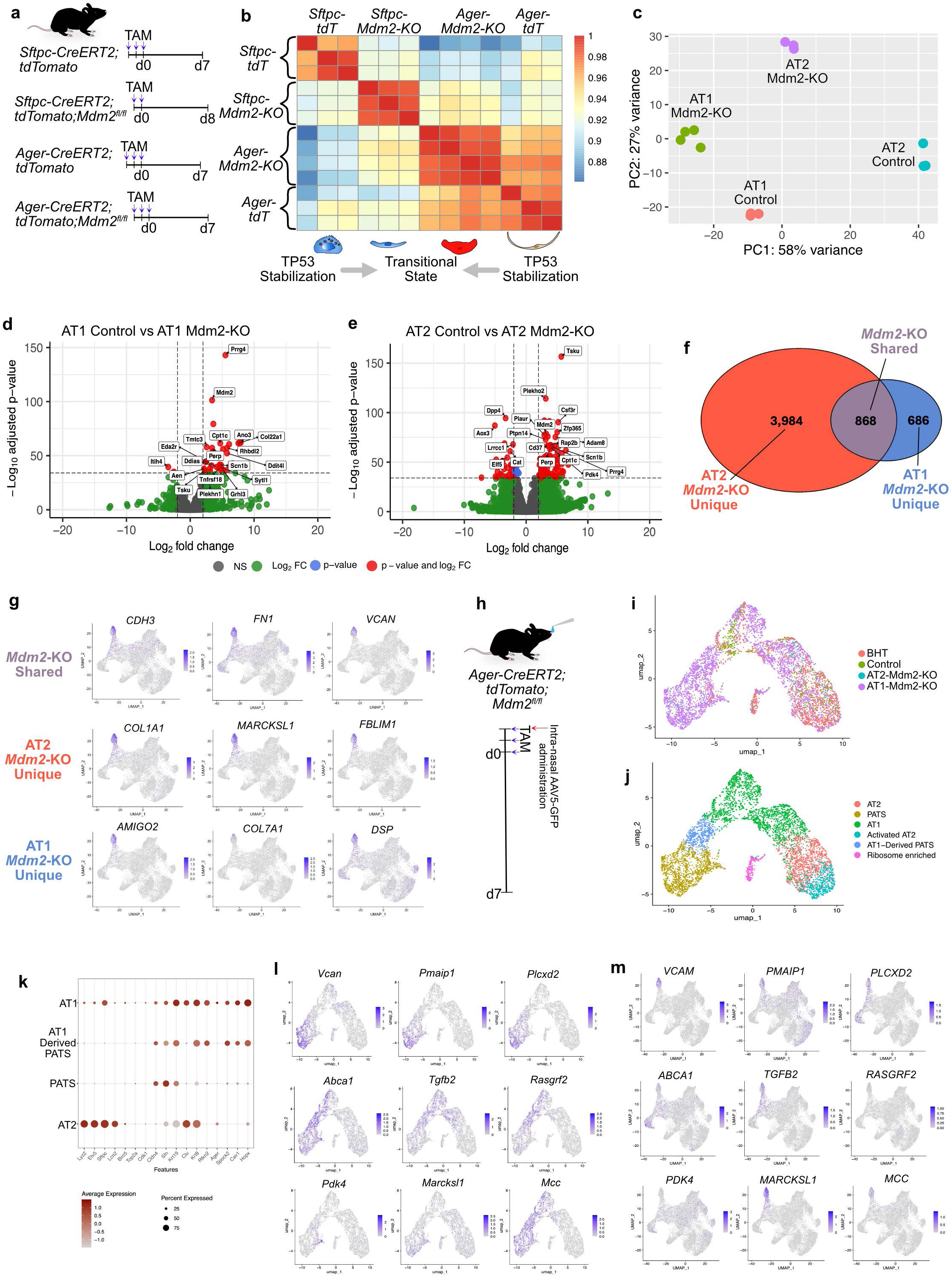
Bulk and single-cell transcriptomics reveal shared and distinct PATS signatures originated from AT2 and AT1. a. Schematic illustrating mice utilized for bulk RNA sequencing, highlighting controls and experimental groups. All mice undergo lineage labeling, and the experimental mice undergo *Mdm2* deletion in AT2- or AT1-specific promoter. At the day indicated, lineage-labeled cells were collected via FACS. b. Bulk RNA sequencing heat-map, showing gene expression comparisons of AT2 control (*Sftpc-CreERT2;tdTomato*), AT2-Mdm2-KO (*Sftpc-CreERT;tdTomato;Mdm2^fl/fl^*), AT1 control (*Ager-CreERT2;tdTomato*), and AT1-Mdm2-KO (*Ager-CreERT2;tdTomato;Mdm2^fl/fl^*). Schematic underneath shows cell types as illustrations and their convergence upon the PATS state in response to *Mdm2* deletion. c. Bulk RNA sequencing PCA plot, showing clustering of indicated groups. d. Volcano plot of differentially expressed genes detected in bulk RNA sequencing, comparing AT1 control, and AT1-Mdm2-KO, highlighting 20 of the top significant hits. e. Volcano plot of differentially expressed genes detected in bulk RNA sequencing, comparing AT2 control, and AT2-Mdm2-KO, highlighting 20 of the top significant hits. f. Venn Diagram showing differentially expressed genes detected in bulk RNA sequencing, highlighting that 3,984 genes are up-regulated in AT2-Mdm2-KO exclusively (differentially expressed in *Sftpc-CreERT2/tdTomato/Mdm2^fl/fl^* over *Sftpc-CreERT2/tdTomato*), 686 genes are up-regulated in AT1-Mdm2-KO exclusively (differentially expressed in *Ager-CreERT2/tdTomato/Mdm2^fl/fl^* over *Ager-CreERT2/tdTomato*), and 868 genes are up-regulated in both AT2- and AT1-Mdm2-KO. g. Differentially expressed genes identified in bulk RNA sequencing from the “Mdm2-KO Shared”, “AT2-Mdm2-KO”, and “AT1-Mdm2-KO” categories are expressed in a human IPF scRNA-seq dataset. h. Experimental workflow for bi-directional lineage labeling in *Ager-Mdm2-KO* mice through tamoxifen injections and intranasal AAV5-GFP administration followed by sample collection on day 7 for scRNA-seq capture and analysis. I. Single cell data integrated with healthy control, injury model, and AT2-Mdm2-KO as a DimPlot of the epithelial cell types with annotations, highlighting contribution of AT1-derived PATS. j. Single cell data integrated with healthy control, injury model, and AT2-Mdm2-KO, represented as a DimPlot split by the assay of origin, highlighting AT1-Mdm2-KO contribution. k. DotPlot showing expression of key marker genes of alveolar epithelial cell types across the indicated genotypes in the single-cell data. l. Mapping differentially expressed genes identified in bulk RNA sequencing from the “Mdm2-KO Shared”, “AT2-Mdm2-KO”, and “AT1-Mdm2-KO” categories onto single cell data from this AT1-Mdm2-KO model, showing expression of these genes across clusters annotated as both “PATS” and “AT1-derived PATS”. m. Mapping indicates differentially expressed genes identified in bulk RNA sequencing from the “Mdm2-KO Shared”, “AT2-Mdm2-KO”, and “AT1-Mdm2-KO” categories onto human scRNA-seq data set.

To dissect which gene programs contribute to the PATS state in a cell-of-origin-specific manner, we partitioned the differentially expressed genes across these four groups into three categories: (1) AT2-*Mdm2*-KO unique - differentially expressed in *Sftpc-Mdm2-KO* compared to *Sftpc-tdT*; (2) AT1-*Mdm2*-KO unique - differentially expressed in *Ager-Mdm2-KO* compared to *Ager-tdT*; and (3) *Mdm2*-KO shared - differentially expressed in both *Sftpc-Mdm2-KO* and *Ager-Mdm2-KO* but not in either control (Figure 5f, Supplementary Data Table 1). Importantly, when projected onto a published single-cell RNA-sequencing dataset of human IPF lungs^30^, genes from all three categories were strongly enriched in the “PATS-like” aberrant basaloid population that is highly enriched in interstitial lung diseases including idiopathic pulmonary fibrosis (Figure 5g). Together, these data suggest that TP53 stabilization in AT1 and AT2 cells, despite their distinct morphological and molecular signatures, drives convergence onto a shared PATS-associated transcriptional state.

To dissect this convergence at single-cell resolution, we next performed scRNA-seq on the *Ager-Mdm2-KO* model. To distinguish PATS derived from AT1 or AT2 cells, we infected mice with AAV5-GFP prior to tamoxifen administration as previously described to lineage label AT2 cells (Figure 5h, Supplementary Figure 5c, 5d). Lungs were harvested at day 7, and immune cells and fibroblasts were partially depleted by magnetic cell separation (MACS) to enrich epithelial cells. We then integrated this dataset with previously acquired single-cell datasets spanning controls, BHT-injured lungs, and the *Sftpc-Mdm2-KO* model representing AT2-derived PATS (Figure 5i). Epithelial cells were identified and subset based on *Epcam* expression (Supplementary Figure 5e). Unsupervised clustering of the epithelial subset yielded 10 distinct clusters, which we annotated based on the expression of canonical alveolar epithelial markers (Figure 5j). Specifically, *Ager*, *Rtkn2*, and *Spock2* marked AT1 cells (Supplementary Figure 5g); *Sftpc*, *Lamp3*, and *Spink5* marked AT2 cells (Supplementary Figure 5h); and *Mdm2*, *Sfn*, *Cldn4*, *Lgals3*, and *Krt8* marked the PATS state (Supplementary Figure 5f), with each cluster displaying clear separation and the expected marker enrichment (Figure 5k). Critically, tdTomato^+^ AT1 lineage-labeled cells extended beyond clusters expressing strictly AT1 markers and spilled into a cluster co-expressing both PATS and AT1 markers (Supplementary Figure 5l). Together, these scRNA-seq data corroborate our observations in human tissue, genetic mouse models, and *in vitro* cultures, indicating that stabilization of TP53 in AT1 cells drives their transition into an intermediate, PATS-like alveolar epithelial state.

Intriguingly, upon mapping AT1-unique, AT2-unique, and *Mdm2*-KO shared gene sets from bulk RNA-seq onto the single-cell data, genes from all three categories including canonical PATS markers occupied PATS clusters (Figure 5l). We also identified a subset of differentially expressed genes that are distinct between AT1- and AT2-specific *Mdm2*-KO cells (Figure 5m). Taken together, these data suggests that certain PATS-associated genes remain preferentially linked to AT1 or AT2 cell-of-origin transcriptional profiles.

### AT1-derived PATS molecular signatures mark regions of alveolar epithelial detachment in human pulmonary disease

Lastly, we sought to leverage our transcriptomic characterization of AT1-derived PATS to define their representation in human tissue and their contribution to disease. To this end, we employed USHER^38^ using the same basaloid gene module score as in Figure 1g as a reference, but additionally interrogated regions of detached AT1-like cells with three new module scores derived from our bulk and single-cell datasets: “Shared PATS”, “AT2-derived PATS”, and, most importantly, “AT1-derived PATS”. As expected, regions exhibiting AT1-like cell detachment scored highly for the basaloid gene module relative to AT1 and AT2 cells (Figure 6a). Notably, the same regions were also enriched for the “Shared PATS” module (*ANXA8*, CDH3, *GDF15*) (Figure 6b) and for the “AT2-derived PATS” module (*ARL4C*, *BACE2*, *CDKN2A*, *CHL1*, *CST6*, *CRLF1*, *KCNN4*, *KRT17*, *LY6D*, *TMEM59L*) (Figure 6c). Critically, regions of detached AT1-like cells were also strongly enriched for the “AT1-derived PATS” module (*COL7A1*, *COL17A1*, *DSP*, *EPHX3*, *LYPD1*, *SYT8*, *WNK2*) (Figure 6d and Supplementary Figure 6a-d). Together, these data indicate that the AT1-derived PATS state we identified in our mouse models is recapitulated in AT1-like cells that detach from the basement membrane in human fibrotic lungs.

**Figure 6:**
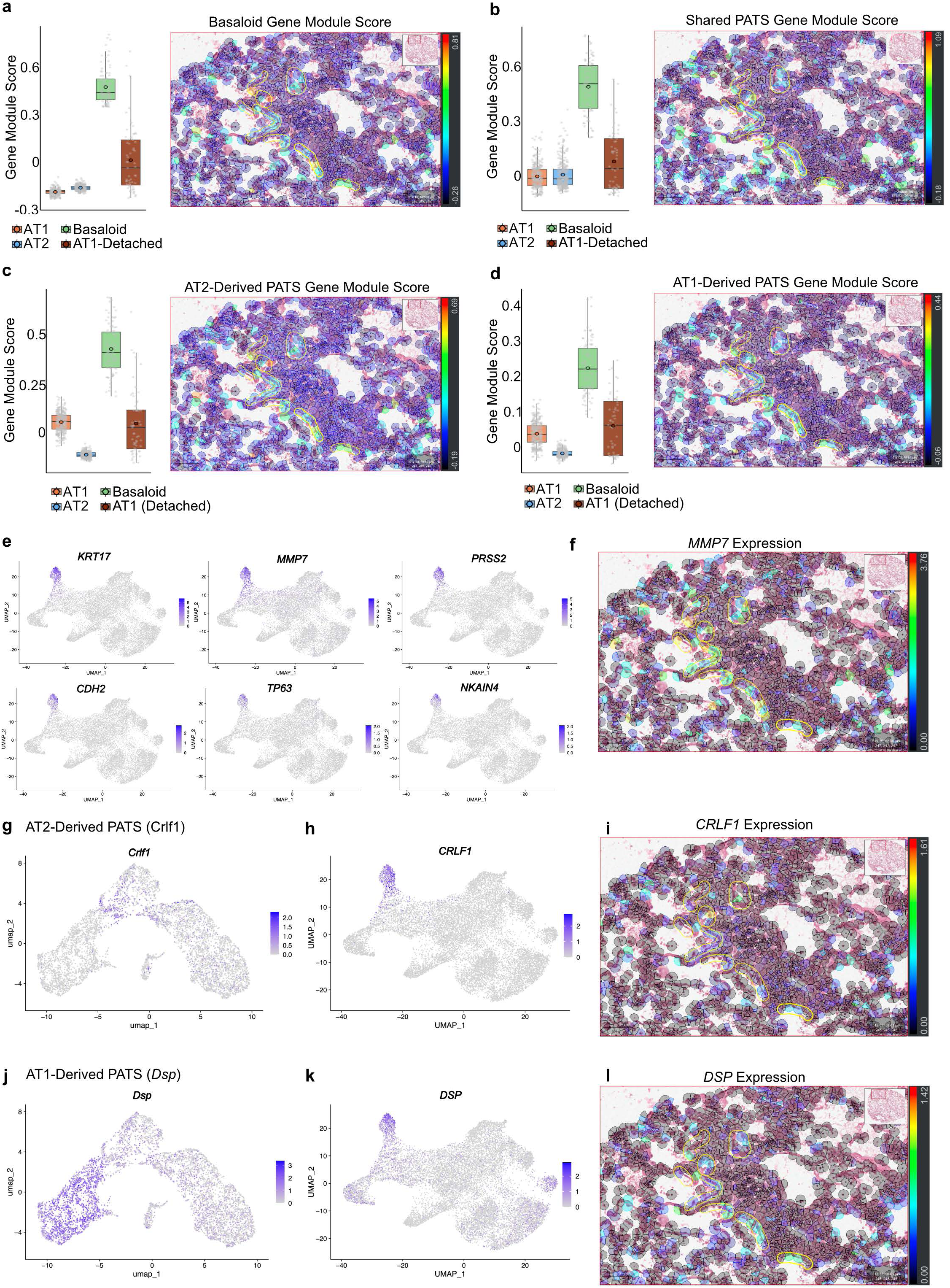
Integrated single cell and spatial transcriptome revealed signatures of AT1-derived PATS in regions of AT1 detachment within diseased human tissue. a. Basaloid gene module score in human tissue, comparing cells classified as AT1, AT2, and basaloids, with regions of AT1 detachment. USHER section shows scoring of individual cells within regions of AT1 detachment, circled in yellow. b. Shared PATS gene module score in human tissue, comparing cells classified as AT1, AT2, and basaloids, with regions of AT1 detachment. USHER section shows scoring of individual cells within regions of AT1 detachment, circled in yellow. c. AT2-derived PATS gene module score in human tissue, comparing cells classified as AT1, AT2, and basaloids, with regions of AT1 detachment. USHER section shows scoring of individual cells within regions of AT1 detachment, circled in yellow. d. AT1-derived PATS gene module score in human tissue, comparing cells classified as AT1, AT2, and basaloids, with regions of AT1 detachment. USHER section shows scoring of individual cells within regions of AT1 detachment, circled in yellow. e. FeaturePlots showing genes from the Basaloid Module Score expression in human scRNA-seq dataset. f. USHER showing expression of *MMP7*, one of the genes in the Basaloid Module Score, within regions of AT1 detachment, circled in yellow. g. FeaturePlot showing expression of *Crlf1*, one of the genes in the AT2-derived PATS Module Score, in our mouse scRNA-seq dataset. h. FeaturePlot showing expression of *CRLF1*, one of the genes in the AT2-derived PATS Module Score, in a human scRNA-seq dataset. i. USHER showing expression of *CRLF1*, one of the genes in the AT2-derived PATS Module Score, within regions of AT1 detachment, circled in yellow. j. FeaturePlot showing expression of *Dsp*, one of the genes in the AT1-derived PATS Module Score, in our mouse scRNA-seq dataset. k. FeaturePlot showing expression of *DSP,* one of the genes in the AT1-derived PATS Module Score, in a human scRNA-seq dataset. l. USHER showing expression of *Dsp*, one of the genes in the AT1-derived PATS Module Score, within regions of AT1 detachment, circled in yellow.

To further validate the molecular concordance between our mouse-derived PATS gene programs and human disease, we systematically cross-referenced each module across mouse scRNA-seq, human scRNA-seq, and USHER imputed spatial transcriptome data. First, we confirmed that genes comprising the basaloid gene module score were robustly represented within the PATS-like cluster of the human scRNA-seq dataset (Figure 6e) and were similarly detected at high levels by USHER, as exemplified by *MMP7* (Figure 6f). Similarly, genes contributing to the “AT2-derived PATS” module were represented in both our mouse (Figure 6g) and the human IPF (Figure 6h) single-cell datasets and were detected at high levels in USHER imputed spatial transcriptome data (Figure 6i), as exemplified by *Crlf1*. Finally, genes contributing to the “AT1-derived PATS” module were likewise represented in mouse (Figure 6j) and human (Figure 6k) single-cell data and were detected at high levels in spatial transcriptome data, as exemplified by *Dsp* (Figure 6l). Taken together, by leveraging USHER imputed spatial transcriptome data alongside mouse and human single-cell transcriptomics, we demonstrate that AT1-derived PATS cells are not restricted to our genetic mouse models but are also evident in human pulmonary disease.

## Discussion

The fitness of a tissue depends critically on the proper functioning of its constituent cells, and dysregulated cellular identity inevitably compromises tissue homeostasis. Defining the limits of cellular plasticity in the alveolar epithelium is therefore central to identifying cell types that can be targeted for the treatment of lung diseases. In this work, we demonstrate that, although AT1 cells possess a largely restricted fate, they can in fact “revert” along the alveolar epithelial continuum and adopt a cell state that converges upon the previously characterized AT2-derived PATS. In recent years, the field has come to recognize PATS as both an obligate transitional state during healthy alveolar repair and as an aberrant, uncontrolled cell state in disease. However, central questions remain unanswered - most notably the etiology of “idiopathic” pulmonary fibrosis and the cellular and molecular drivers of its progression. While ongoing work continues to refine our understanding of AT2-derived PATS, AT1 cells have largely been treated as terminally differentiated cells that contribute to disease solely through their loss. Our lineage-tracing of AT1-derived PATS now reveals that, although AT1 cell loss is itself a crucial contributor to alveolar remodeling, AT1-derived PATS cells express many of the same genes that define the “pro-fibrotic” character of AT2-derived PATS. Notably, these AT1-derived PATS cells are themselves susceptible to extrusion, and their loss from the alveolar epithelium further mobilizes a subsequent AT2-derived PATS response. Together, these findings suggest a “bidirectional” induction of PATS that can establish a runaway pathological cycle: AT1-derived PATS loss elicits AT2-derived PATS, which remodel the alveolar architecture and compress neighboring AT1 cells, in turn inducing further AT1-derived PATS and perpetuating this pattern.

Critically, our work demonstrates that AT1-derived PATS can be acquired through cell-intrinsic mechanisms in the complete absence of physical injury. This is of particular relevance because age is among the strongest risk factors for IPF, and TP53 signaling is known to accumulate in aged lung tissue. Additionally, TP53 signaling has been shown to play a role in cell extrusion^41–43^ in cancer tissues and we propose that TP53 does so via induction of transitional state and driving their loss via cell extrusion.

An important open question that deserves dedicated attention in future studies is how extrinsic cues influence this phenomenon specifically, whether mesenchymal remodeling in the form of interstitial thickening and myofibroblast-mediated constriction can in turn stabilize TP53 in adjacent AT1 cells and thereby induce an AT1-derived PATS state. It has been demonstrated that TP53 is necessary and sufficient for cells to become hypersensitized to crowding^41,44^ and that crowding contributes directly to extrusion^2,3,37^. Defining the interplay between such cell-intrinsic and microenvironmental drivers will be essential to understanding how the bidirectional PATS cycle is initiated and sustained in human disease.

Additionally, open questions remain as to mechanistically how TP53 stabilization in AT1 cells induces this transitional state and eventual extrusion. Previous work has shown that TP53 activation is involved in the synthesis and turnover of sphingosines^45^, and sphingosine signaling is upstream of Rho-mediated actomyosin contraction that is necessary for successful cell extrusion^37^. Additionally, it has been demonstrated that replication stress and subsequent TP53 activity contribute to extrusion^46^, as well as mitochondrial stress^47,48^. Interestingly, in the case of mitochondrial stress, it was observed that AT1 cells undergoing mitochondrial stress were “thicker and rounder” than control AT1 cells, SFTPC-expressing cells “exhibited a linear and thin shape more typical of AT1 cells”^48^, which aligns with cellular identity dysregulation observed in *Ager-Mdm2-KO* lungs. Critically, previous work suggested that mitochondrial stress in alveolar epithelial cells renders them for enhanced liberation of rounder and less mature cells”^48^, implying increased vulnerability to extrusion, as we observed in our experiments. Similarly, pyruvate dehydrogenase kinase 4 (PDK4) has been shown to drive mitochondrial dysfunction and regulated cellular extrusion^47^. This is in line with our scRNA-seq data that uncovered *Pdk4* as one of the top markers within a unique cluster of AT1-derived PATS, which likely represents extruding cells.

Together, these findings provide a new conceptual framework for understanding human IPF. The contribution of dysregulated AT1-derived PATS aligns well with the cardinal features of the disease, namely, a paucity of AT1-containing gas-exchange regions, the accumulation of ectopic intermediate alveolar epithelial cells, and persistent myofibroblast activation. We anticipate that this new perspective will guide future investigations into previously overlooked therapeutic approaches that target the AT1-derived PATS state itself, rather than focusing solely on AT2 cell dysregulation, opening new avenues for the treatment of IPF and related fibrotic lung diseases.

## Materials and Methods

### EXPERIMENTAL MODEL AND SUBJECT DETAILS

#### Animals

*S*ftpc*^tm1(cre/ERT2)Blh^ (Sftpc-CreER)* (stock number 028054, Jackson Laboratory), *Rosa26R-CAG-lsl-tdTomato (Ai14)* (stock number 007914, Jackson Laboratory), *Ager^tm1(cre/ERT2)Blh^ (Ager-CreER)* (stock number 036942, Jackson Laboratory), *Mdm2^tm2.1Glo^*/J (stock number 031614, Jackson Laboratory), and C57BL/6J (stock number 000664, Jackson Laboratory) mice were maintained on a C57BL/6J background. For all lineage tracing or loss of function experiments using *Ager-CreERT2* driver, 3 doses of tamoxifen (0.2 mg/g body weight) were used. For bulk RNA sequencing experiments, *Sftpc-CreERT2* driver line mice received 2 doses of tamoxifen for controls and 2 doses of tamoxifen for Mdm2-KO (0.1 mg/g body weight). AAV5-GFP was intranasally administered to mice as described previously^49^. Both male and female mice between 8 and 16 weeks of age were used for experiments. All animal experiments were approved by the Duke University Institutional Animal Care and Use Committee in accordance with US National Institutes of Health guidelines.

### METHOD DETAILS

#### Human Immunofluorescence staining

Paraffin sections were heated to 65°C and subsequently incubated in Histo-Clear, 100% EtOH, 75% EtoH, 50% EtOH, and water, for 5 minutes each to dewax and rehydrate the tissue. Antigen retrieval was performed in EDTA buffer, pH 8.5 (Sigma-Aldrich) or citrate buffer, pH 6.0 (Sigma-Aldrich) using a water bath (95°C for 10 min). Sections were washed with 0.1% Triton in PBS (PBST), incubated in blocking buffer (1% BSA in 0.1% PBST) for 1 h, and then stained with primary antibody in blocking buffer overnight at 4°C. Following primary antibody incubation, autofluorescence was quenched using the TrueVIEW autofluorescence quenching kit (Vector labs, SP-8500) and tissue was washed three times in PBST, followed by incubation with secondary antibody in blocking buffer for 1h at RT. Sections were washed with PBST and coverslips were mounted with Fluor G reagent with DAPI. Primary antibodies were as follows: LEL-DyLight-488 (Vector Laboratories, DL-1174-1) at 1:1000, Nkx2-1 (Abcam, ab133737) at 1:500, HTII-280 (Terrace Biotech, TB-27AHT2-280) at 1:500, p53 (Alexa Fluor® 594; BioLegend, 645706) at 1:250, and SFN (Proteintech, 66251-1-Ig) at 1:500.

#### Mouse Tissue preparation and sectioning

Lungs were inflated with 4% Paraformaldehyde (PFA) and incubated at 4°C for 4–6 h. Lung lobes were separated, washed in PBS, and incubated overnight in 30% sucrose at 4°C. Lobes were subsequently incubated in 1:1 30% sucrose:OCT for 1 h followed by embedding in OCT blocks and cryosectioning. For all 2D imaging, sections were collected at 8µm. For thick sections, lungs were inflated with 1.5mL low melting point 1.5% agarose dissolved in 1x PBS. Lungs were placed on ice until agarose solidified, followed by vibratome sectioning at 750µm. PCLS sections were either live-imaged or collected in PFA, fixed for 4h at 4°C and stored in PBS until further processing.

#### Mouse Immunofluorescence staining

OCT sections were brought to room temperature and washed in PBS. Antigen retrieval was performed in EDTA buffer, pH 8.5 (Sigma-Aldrich) or citrate buffer, pH 6.0 (Sigma-Aldrich) using a water bath (95°C for 10 min). Sections were washed with 0.1% Triton in PBS (PBST), incubated in blocking buffer (1% BSA in 0.1% PBST) for 1 h, and then stained with primary antibody in blocking buffer overnight at 4°C. Following primary antibody incubation, tissues were washed three times in PBST followed by incubation with secondary antibody in blocking buffer for 1 h. Sections were washed with PBST and coverslips were mounted with Fluor G reagent with DAPI. Primary antibodies were as follows: LEL-DyLight-488 (Vector Laboratories, DL-1174-1) at 1:1000, LEL-DyLight-649 (Vector Laboratories, DL-1178-1) at 1:1000, p53 (Alexa Fluor® 647; Santa Cruz, sc-126 AF647) at 1:250, SFN (Proteintech, 66251-1-Ig) at 1:500, HOPX (Millipore, HPA030180) at 1:250, TdTomato (Origene, AB8181-200) at 1:700, LAMP3 (Synaptic Systems, 391 005) at 1:1000, GFP (Novus Biologicals, NB100-1770) at 1:500, Sftpc (Proteintech, 10774-1-AP) at 1:1000, MAC2 (Galectin-3; Cedarlane, CL8942AP) at 1:500, Ki67 (Invitrogen, 14-5698-82) at 1:250, and αSMA (Abcam, ab5694) at 1:250.

#### Image acquisition and processing

Images were captured on an Olympus FV3000 confocal microscope using 20X, 40X, and 60x objectives. Images from lung tissue staining are presented as a z-projection over approximately 2-4 µm unless otherwise stated.

#### Live imaging

750µm PCLS sections were incubated in Advanced DMEM with LEL-DyLight-649 for 30 minutes at 37°C. Sections were submerged in Advanced DMEM and imaged for 225 frames at 4-minute intervals. Recordings were processed in Imaris software. Cell masks were applied using automated selection criteria. Representative still frames are taken from recordings.

#### BAL collection

Mice were processed as previously described, except rather than inflating lungs with 4% PFA, 1 mL of 1x PBS was injected into the lungs and aspirated repeatedly for a total of 3 times. Lavage fluid was then placed in a 15 mL tube, and repeated 3 times, for a total volume of 3 mL. This suspension was centrifuged at 500 x g at 4°C. The cell pellet was fixed in 200 µL of 4% PFA at RT for 15 minutes. Cells were once again centrifuged at 500 x g at 4°C and resuspended in 200 µL PBS. 100 µL of cell suspension was spun onto slides and mounted with Fluoromount-G with DAPI for imaging.

#### AT1 Cultures

*Ager-CreERT2/tdTomato/MDM2^fl/fl^* mouse AT2s were isolated via FACS as previously described^50,51^ and induced to differentiate into AT1s as previously described^39^. Fibronectin (Santa Cruz, SC-29011A) was diluted with PBS to a concentration of 50 μg/mL, then added to wells at room temperature for 30 min. Fibronectin was removed, and wells were washed once with PBS followed by AT2 seeding diluted in culture medium. The media was changed every three days. At day 9, when AT2s were differentiated into AT1s, Adeno-Cre virus was added to the media to induce Cre-mediated deletion of *Mdm2* and expression of tdTomato in AT1s. Adeno-GFP virus was used as control.

#### Mouse lung tissue dissociation and cell isolation

Lung dissociation was performed as described previously^50^. Briefly, lungs were inflated with an enzyme dissociation solution (450U/mL Collagenase I (Worthington, LS004197), 5U/mL Dispase (Corning, 354235), and 0.33U/mL DNase I (Roche, 10104159001), in DMEM). Separated lung lobes were minced and incubated in enzyme solution at 37°C for 25–35 min. Dissociation was quenched with equivalent volume of 10% FBS/DMEM and strained through a 100µm strainer. Cell pellet was resuspended in red blood cell lysis buffer (100 mM EDTA, 10 mM KHCO3, 155 mM NH4Cl) for 2 min, followed by quenching with 10% FBS/DMEM and filtration through a 40µm strainer.

#### RNA preparation for bulk RNA-seq

tdTomato-positive cells were isolated by fluorescence-activated cell sorting (FACS) from the lungs of *Ager-CreERT2/tdTomato/Mdm2^fl/fl^* experimental (n=4), *Ager-CreERT2/tdTomato* control (n=3), *Sftpc-CreERT2/tdTomato/Mdm2^fl/fl^* experimental (n=3), and *Sftpc-CreERT2/tdTomato* control (n=3) mice. Sorted cells were resuspended in TRIzol reagent (Thermo Fisher Scientific, 15596026) and total RNA was extracted using a Direct-zol RNA Microprep kit (Zymo, R2061) with DNase I treatment according to the manufacturer’s protocol.

#### Bulk RNA sequencing and differential gene expression analysis

Ribosomal RNA was depleted from total RNA samples (100 ng) using NEBNext rRNA Depletion Kit v2 (New England BioLabs, E7400L). Libraries were prepared using NEBNext Ultra II Directional RNA Library Prep Kit for Illumina (New England BioLabs, E7760S). Paired-end sequencing (150 bp for each read) was performed on an Illumina platform. Raw sequencing reads were quality-filtered using FaQCs^52^ with adapter trimming, poly-A removal, and quality thresholds (Q30 average, Q10 minimum, 15 bp minimum length). Trimmed reads were aligned to the mouse reference genome (UCSC mm10) using STAR^53^ with two-pass mode. Transcript assembly and abundance estimation were performed using StringTie^54^ in reference-guided mode, and gene-level expression matrices were generated using tximport^55^. Normalization and extraction of differentially expressed genes (DEGs) between samples were performed using an R package, DESeq2^56^. Differentially expressed genes were defined as log2FC x>1 or x<-1 with p-adj x<0.05.

#### Single-cell RNA sequencing

n=2 *Ager-CreERT2/tdTomato/Mdm2^fl/fl^* experimental mice were harvested for lung tissue. MACS based immune and endothelial cell depletion was performed to enrich epithelial and mesenchymal cells. Cells from dissociated lungs were incubated for 30 min with the following antibodies: CD45 (MiltenyiBiotech, 130052-301) and CD31(MiltenyiBiotech,130-097-418). MACS depletion was performed as per manufacturer protocol using LD columns (MiltenyiBiotech, 130-042-901). After cell purification, 10% spike-in of total cells were added to each sample, and the samples were mixed evenly. We generated a single cell transcriptome library targeting 20,000 cells using the 10X Genomics GEMX 3’ Capture v4 kit (10X Genomics PN-1000691). The sample was sequenced using the Novaseq X platform with a sequencing depth of 400M paired reads.

#### Analysis of single cell transcriptomic libraries

To process and analyze the sample generated in this study, we utilized Cellranger v9^57^ and Seurat v4^58^. Sample fastq files were processed with Cellranger v9.0.1 to align to the mm10 genome. Raw count matrices from the *Ager-Mdm2-KO* sample and previously processed Control, BHT, and *Sftpc-Mdm2-KO* mouse lung datasets were extracted^40^ and reconstructed as independent Seurat objects. Doublets were identified per sample using DoubletFinder^59^ following LogNormalize preprocessing, PCA (30 PCs), and clustering. The optimal pK for each sample was selected by maximizing the BCmvn metric, and the expected doublet rate was set to 0.8% per 1,000 cells. Cells classified as doublets were removed. Singlet count matrices were normalized per sample using SCTransform and integrated using FindIntegrationAnchors and IntegrateData with SCT normalization. The integrated assay was scaled, and PCA, UMAP, and Louvain clustering were performed using the first 12 principal components (resolution = 1.0). Non-epithelial and contaminating populations were removed iteratively based on lineage marker expression (*Epcam*, *Pdgfra*, *Ptprc*, *Pecam1*, *Sftpc*), and cells were further filtered to retain those with ≤ 5% mitochondrial reads and 500–6,000 detected features. Cluster markers were identified using FindAllMarkers on the RNA assay (only.pos = TRUE, min.pct = 0.25, logfc.threshold = 0.25).

#### USHER Analysis

All analysis was done within QuPath-0.5.1. H&E images were overlaid with spatial transcriptomics data and imputed single cell data. Measurements were made as both single gene scores as well as gene module scores, comprised of multiple genes as follows: AT1 (*SPOCK2*, *TIMP3*, *AGER*, *RTKN2*, *CAV1*, *UPK3B*, *RGCC*, *WFS1*, *COL4A2*, *NCKAP5*, *MS4A15*, *FMO2*, *COL12A1*, *MYRF*, *PLAC8*, *SLC1A1*, *ITLN2*, *UNC13D*, *COL4A1*, *ANKRD1*), AT2 (*SFTPC*, *SFTPA1*, *PGC*, *LRRK2*, *LAMP3*, *ABCA3*, *HHIP*, *TTN*, *ALOX15B*, *CA2*, *HP*, *C4BPA*, *PLA2G1B*, *NRGN*, *ETV1*, *SLC22A31*, *CACNA2D2*, *KIAA1324L*), Basaloid (*KRT17*, *MMP7*, *PRSS2*, *PLAU*, *CDKN2A*, *CDH2*, *KCNN4*, *TP63*, *NKAIN4*, *AC025580.1*), TP53(*SFN*, *GDF15*, *BAX*, *BBC3*, and *RRM2B*), Shared PATS (*ANXA8*, *ARNTL2*, *BBC3*, *CAPG*, *CDH3*, *EDA2R*, *ESPN*, *FAS*, *FAT1*, *FHL2*, *FN1*, *GDF15*, *PROCR*, *TGFBI*, *WNT10A*), AT2-Derived PATS (*ARL4C*, *BACE2*, *CDKN2A*, *CHL1*, *CST6*, *CRLF1*, *KCNN4*, *KRT17*, *LY6D*, *TMEM59L*), AT1-Derived PATS (*COL7A1*, *COL17A1*, *DSP*, *EPHX3*, *LYPD1*, *SYT8*, *WNK2*).

Regions of interest were identified as regions of cells that looked like AT1-morphologically but were visibly detaching from the basement membrane. These regions were selected and the cells within these hierarchies were annotated as “detached AT1-like cells”, for a total of 59 cells. AT2 and AT1 cells were selected utilizing a classifier based on gene module score, and 300 cells with the top score from each were selected as controls. 88 cells were classified as Basaloid based on the gene module score.

#### Statistical analysis

Sample size was not pre-determined. All experiments were performed on at least three biological replicates (except single-cell RNA sequencing, which consisted of two biological replicates). Data are presented as means with standard error (SEM). Statistical analysis was performed in Excel, GraphPad Prism, and R. A two-tailed Student’s t-test was used for all comparisons between two conditions. Statistical significance was set a priori at p < 0.05. All data are presented as mean ± SEM. Statistical analyses were performed in R (version 4.5.1). All graphs were generated using ggplot2 (ggplot2_4.0.2).

#### Data quantification

All image-based quantifications were performed in FIJI (ImageJ2). Confocal images were acquired on an Olympus FV3000 with a 20× objective. For every analysis below, data were first tested for normality (Shapiro-Wilk) and homogeneity of variance (Levene’s test) to guide selection of the appropriate statistical test, results are presented with individual biological replicates overlaid, and significance thresholds are defined as *p < 0.05, **p < 0.01, and ***p < 0.001.

#### Lineage-labeled AT1 cells expressing AT2 markers (DCLAMP, SFTPC)

tdTomato⁺ nuclei and tdTomato⁺/DCLAMP⁺ or tdTomato⁺/SFTPC⁺ double-positive nuclei were counted from merged-channel images, and the percentage of lineage-labeled AT1 cells expressing each AT2 marker was calculated as double-positive nuclei divided by total tdTomato⁺ cells. Normality and equal-variance assumptions were satisfied, so group differences were assessed by one-way ANOVA followed by Dunnett’s post-hoc test against the D7 control. Boxplots.

#### tdTomato⁺ cells in bronchoalveolar lavage (BAL)

Total and tdTomato⁺ nuclei were counted and expressed as the tdTomato⁺ fraction. The percentage data satisfied normality and equal-variance assumptions and were compared by Welch’s two-sample t-test; total cell number did not meet normality assumptions and was compared by Mann-Whitney U (Wilcoxon rank-sum) test. Boxplots.

#### AT1 cell size in 2D culture

tdTomato⁺ (Adeno-Cre) and GFP⁺ (Adeno-GFP) cell boundaries were manually traced from the corresponding fluorescence channels using the Freehand Selection tool at days 2 and 3 of culture, and cell area was calculated with the Measure function. Because the small sample size (n = 3 biological replicates per group) limits the power of Shapiro-Wilk to detect deviations from normality, and the design was matched (paired Adeno-GFP vs Adeno-Cre wells from the same culture), comparisons at each timepoint were performed using a paired two-sample t-test. Bar graphs with mean ± SD.

#### ACTA2⁺ tissue coverage

Images were converted to 8-bit, thresholded at 5-255, and signal-positive pixels were extracted from the histogram; tissue coverage was calculated as signal-positive pixels divided by total pixels. Normality (p = 0.30) and equal-variance (p = 0.31) assumptions were satisfied, and conditions were compared by Welch’s two-sample t-test. Boxplots.

## Data Availability

Bulk RNA-seq and single-cell RNA-seq data generated in this study have been deposited in the NCBI Gene Expression Omnibus (GEO) and are publicly available under accession numbers GSE335749 and GSE335750, respectively.

## Acknowledgements

We thank Tata lab members for fruitful discussions, and the Duke Compute Cluster for server space and data storage. We acknowledge and thank the BioRepository and Precision Pathology Center and Research Support-Duke Surgery for providing human tissues under Institutional Review Board oversight and Substrate Services Core.

## Funding support

This work is the result of NIH funding, in whole or in part, and is subject to the NIH Public Access Policy. Through acceptance of this federal funding, the NIH has been given a right to make the work publicly available in PubMed Central. Flow Cytometry core facility is supported by the NCI Cancer Center Support Grant (P30CA014236). T.W is supported by Duke Program of Training in Pulmonary Research to Promote, Engage and Retain Academic Researchers (PROSPER) T32 - 5T32HL160494-05. J.M. was supported by Developmental and Stem Cell Biology Program T32 - T32-HD040372 and Ruth L. Kirschstein National Research Service Award (NRSA) (5F31HL172360) from NIH. This work was supported by research award from NHLBI/NIH (R01HL146557, R01HL160939, R01HL153375 to P.R.T and R01HL174525 to A.T).

## Author Contributions

J.M. led the study, co-designed and performing experiments, analyzed data, and co-wrote the manuscript. K.E. performed ex vivo experiments and analyzed data. T.W. and N.M. generated RNA-seq libraries and performed transcriptomic data analysis. A.P. developed the USHER computational pipeline and contributed to spatial transcriptomic analysis. A.T. co-designed and supervised the study and performed image acquisition. R.S. supervised the work. P.R.T. conceived and co-designed the study, supervised the work, and co-wrote the manuscript with J.M. All authors reviewed and edited the manuscript.

## Competing Interests

The authors have declared that no conflict of interest exists.

## Supplementary Data Table 1

Differentially expressed genes from Bulk RNA sequencing experiments, with genes listed within each tab representing the categories visualized in the Venn Diagram in Figure 5F: (1) AT2-*Mdm2*-KO unique; (2) AT1-*Mdm2*-KO unique; and (3) *Mdm2*-KO shared. 3,984 genes are up-regulated in AT2-Mdm2-KO exclusively (differentially expressed in *Sftpc-Mdm2-KO* over *Sftpc-tdT*), 686 genes are up-regulated in AT1-*Mdm2*-KO exclusively (differentially expressed in *Ager-Mdm2-KO* over *Ager-tdT*), and 868 genes are up-regulated in both AT2- and AT1-Mdm2-KO. These genes were determined by DESeq2 analysis, with the parameters of log_2_ Fold Change x>1 or x<-1 and adjusted p-value x<0.05.

## Supplementary Video 1

Live imaging of 750µm PCLS processed from *Ager-Mdm2-KO* mouse lungs, with lineage-labeled tdT^+^ cells in red and LEL-649 staining in white. Each frame is a z-stack of 25 slices of 3 µm each, with a frame taken every 4 minutes over the course of 8 hours, using a 20x objective.

## Supplementary Video 2

Live imaging of 750µm PCLS processed from *Ager-tdT* mouse lungs, with lineage-labeled tdT^+^ cells in red and LEL-649 staining in white. Each frame is a z-stack of 25 slices of 3 µm each, with a frame taken every 4 minutes over the course of 8 hours, using a 20x objective.

---

**Supplementary Figure 1:**
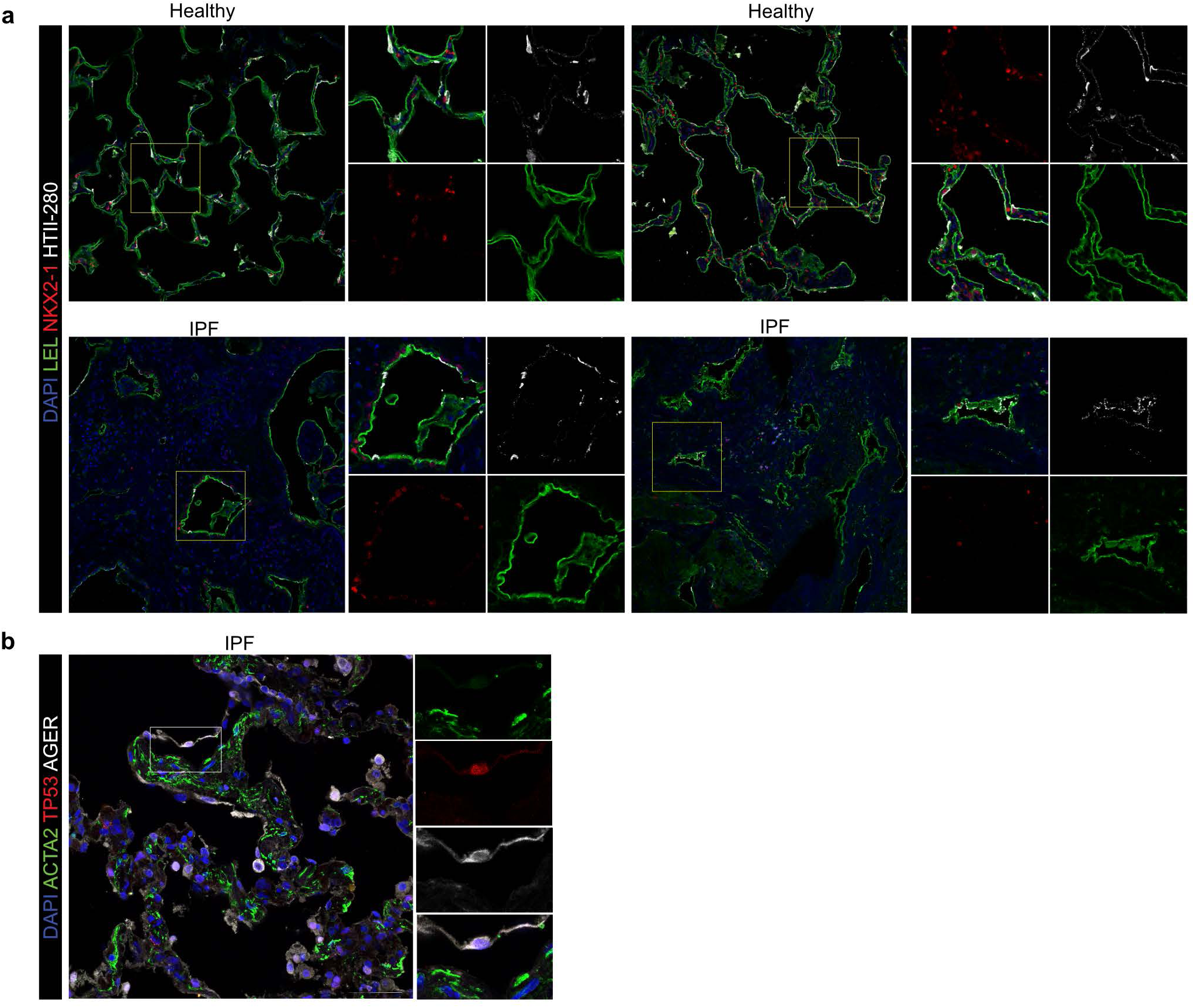
a. Immunostaining for LEL, NKX2-1, and HTII-280 in healthy and IPF human lung tissue. DAPI stains nuclei. Yellow box indicates region of single-channel images. Scale bar: 100 µm. b. Immunostaining for ACTA2, TP53, and AGER in IPF lung. White box indicates region of single-channel images. Scale bar: 100 µm. Images presented as z-projection of 2 stacks over 2 µm.

**Supplementary Figure 2:**
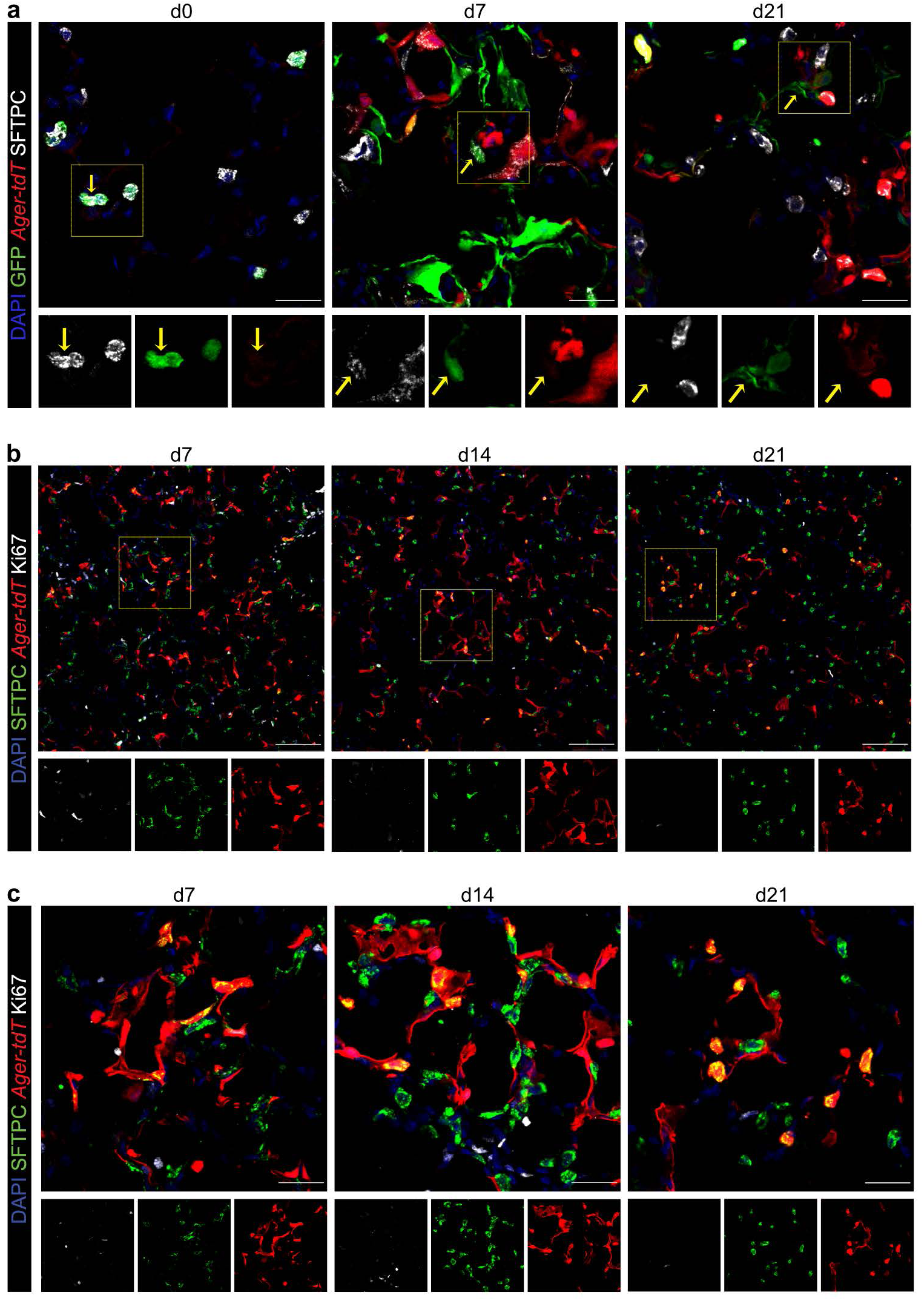
a. Immunostaining for GFP, Ager-tdT, and SFTPC in bi-directional lineage label model at days 0, 7, and 21. Scale bar: 25 µm. b. Immunostaining (low magnification) for SFTPC, Ager-tdT, and Ki67 in *Ager-Mdm2-KO* model at days 7, 14, and 21. Scale bar = 100 µm.. c. Immunostaining (high magnification) for SFTPC, Ager-tdT, and Ki67 in *Ager-Mdm2-KO* model at days 7, 14, and 21. Scale bar = 25 µm. a-c: DAPI stains nuclei. a,b White box indicates region of single-channel images.

**Supplementary Figure 3:**
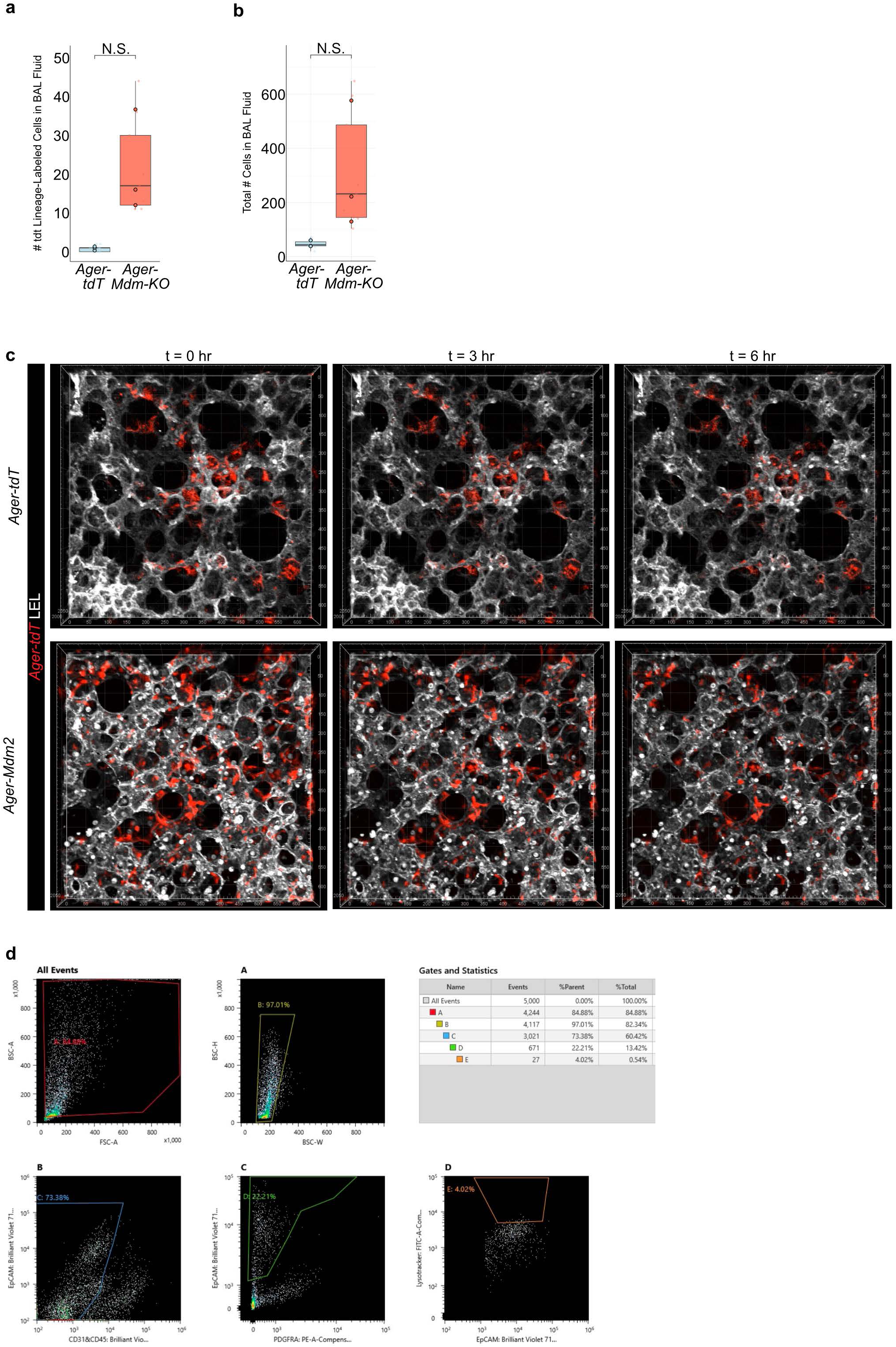
a. Quantification of the number of tdT^+^ lineage-labeled cells per field of view in BAL fluid. Larger opaque points represent biological replicates, smaller transparent points represent technical replicates. n=3. Unpaired two-tailed Welch’s t-test. N.S. p = 0.11. b. Quantification of the total number of cells per field of view in BAL fluid. Larger opaque points represent biological replicates, smaller transparent points represent technical replicates. n=3. Mann-Whitney U test (Wilcoxon rank-sum test). N.S. p = 0.1. c. Time frames from *Ager-tdT* control and *Ager-Mdm2-KO* experimental PCLS cultures, stained for tdT and LEL. Imaris software visualization of full thickness, 25 planes of 3 µm z-projections for a total of 75 µm thickness. Still frames at t=0 hr, t=3 hr, and t=6 hr timepoints. d. FACS gating strategy to isolate AT2 cells from *Ager-Mdm2-KO* experimental mice for 2D culture experiments.

**Supplementary Figure 4:**
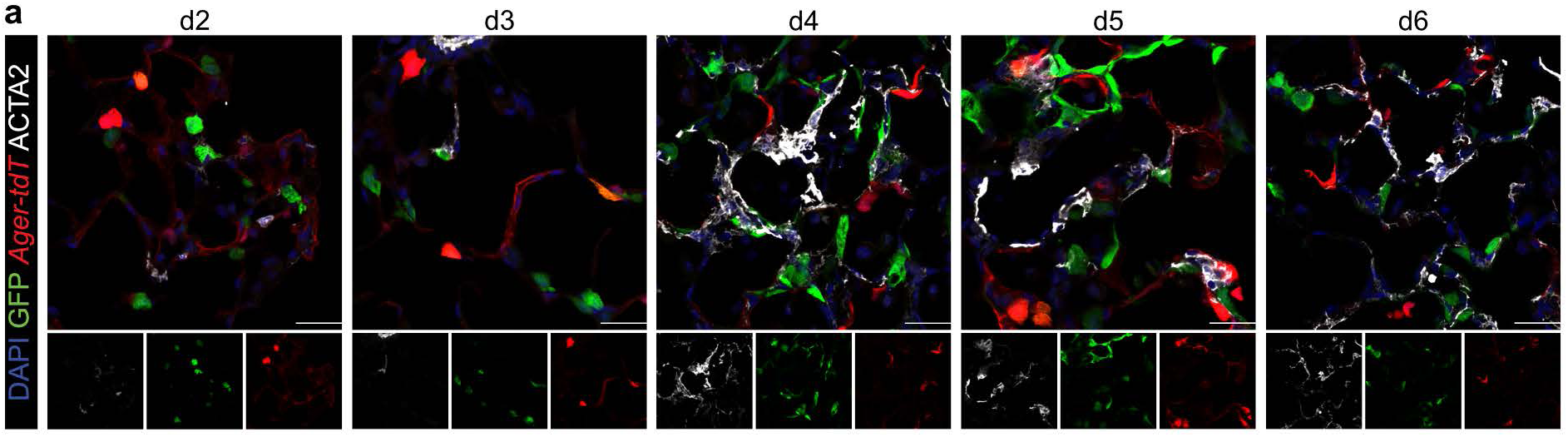
a. Immunostaining for GFP, Ager-tdT, and ACTA2 in bi-directional lineage label model at days 2, 3, 4, 5, and 6. Scale bar: 25 µm.

**Supplementary Figure 5:**
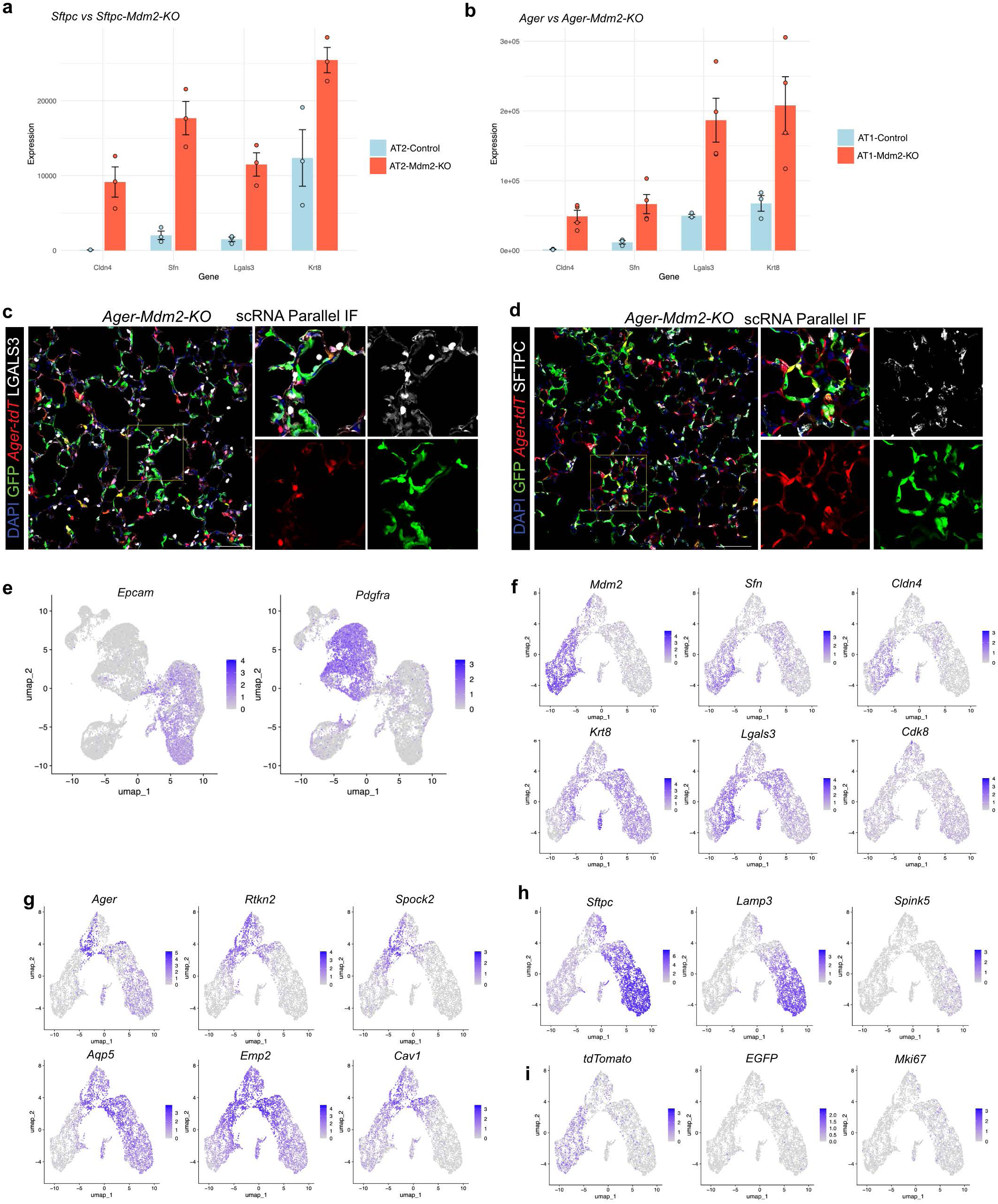
a. Bulk RNA expression of PATS signature genes, *Cldn4, Sfn, Lgals3*, and *Krt8*, in *Sftpc-tdT* compared with *Sftpc-Mdm2-KO*. Each dot represents measure of biological replicate, with error bars representing SEM. b. Bulk RNA expression of PATS signature genes, *Cldn4, Sfn, Lgals3, and Krt8*, in *Ager-tdT* compared with *Ager-Mdm2-KO*. Each dot represents measure of biological replicate, with error bars representing SEM. c. scRNA-seq mouse collected in parallel for IF. Immunostaining for GFP, Ager-tdT, and LGALS3 in bi-directional lineage label model at day 7. Scale bar: 100 µm. d. scRNA-seq mouse collected in parallel for IF. Immunostaining for GFP, Ager-tdT, and SFTPC in bi-directional lineage label model at day 7. Scale bar = 100 µm. e. FeaturePlot from integrated scRNA-seq Seurat object showing expression of epithelial cells (*Epcam*) and mesenchymal cells (*Pdgrfa*) before subsetting epithelial cells for further analysis. f. Feature plots showing expression of PATS identity marker genes in mouse scRNA-seq data, highlighting *Mdm2, Sfn, Cldn4, Krt8, Lgals3,* and *Cdk8*. g. Feature plots showing expression of AT1 identity marker genes in mouse scRNA-seq data, highlighting *Ager, Rtkn2, Spock2, Aqp5, Emp2*, and *Cav1*. h. Feature plots showing expression of AT2 identity marker genes in mouse scRNA-seq data, highlighting *Sftpc, Lamp3*, and *Spink5*. i. Feature plots showing expression of identity marker genes of interest in mouse scRNA-seq data, highlighting *tdTomato, EGFP*, and *Mki67*.

**Supplementary Figure 6:**
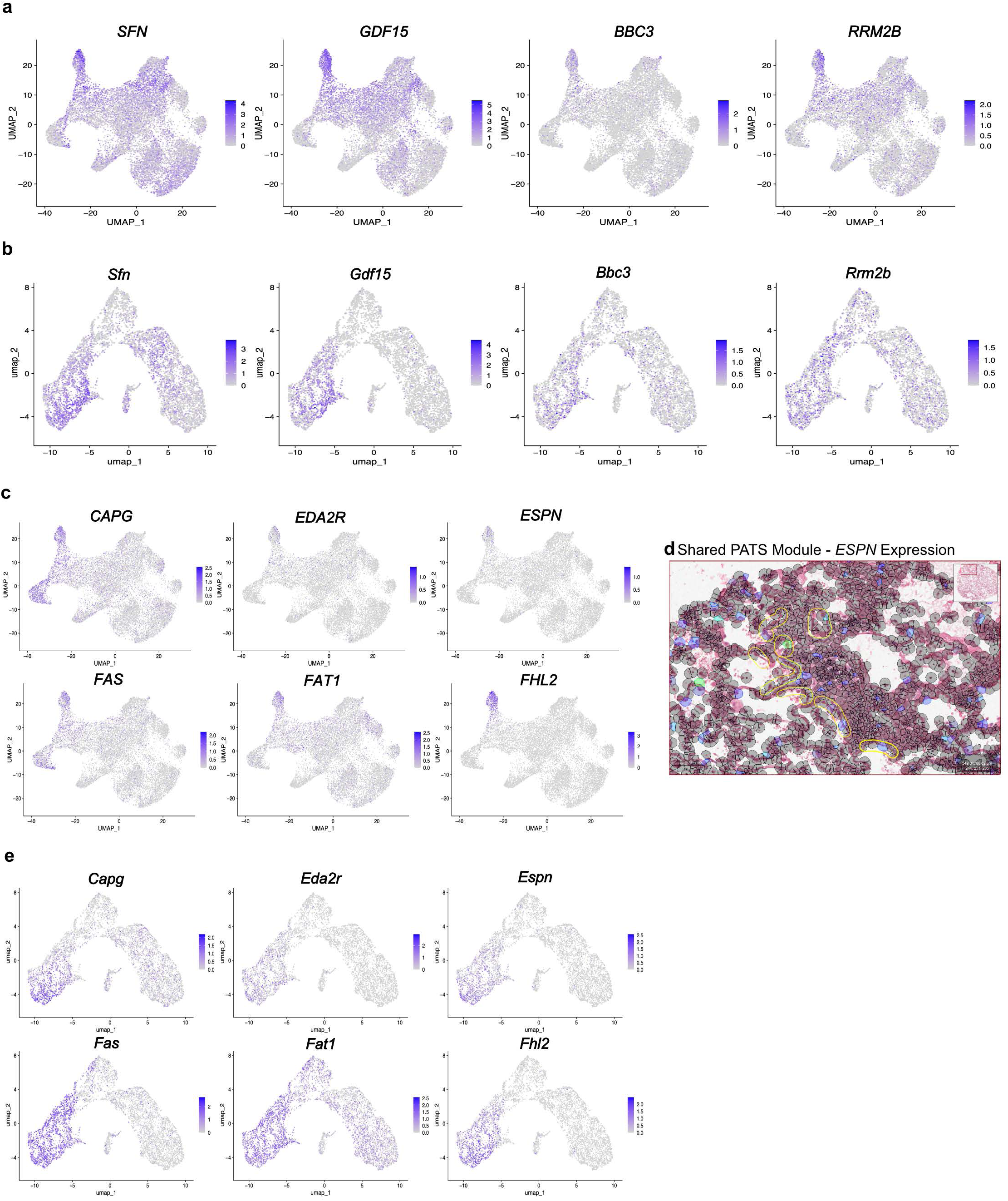
a. Feature plots showing expression of TP53 Signaling Module genes in human scRNA-seq data, highlighting *SFN, GDF15, BBC3*, and *RRM2B*. b. Feature plots showing expression of TP53 Signaling Module genes in mouse scRNA-seq data, highlighting *Sfn, Gdf15, Bbc3*, and *Rrm2b*. c. Feature plots showing expression of Shared PATS Module genes in human scRNA-seq data, highlighting *CAPG, EDA2R, ESPN, FAS, FAT1*, and *FHL2*. d. USHER showing expression of *ESPN*, one of the genes in the shared PATS Module Score, within regions of AT1 detachment, circled in yellow e. Feature plots showing expression of shared PATS Module genes in mouse scRNA-seq data, highlighting *Capg, Eda2r, Espn, Fas, Fat1*, and *Fhl2*.

## Notes

### Competing Interest Statement

The authors have declared no competing interest.

